# Single molecule spectrum dynamics imaging with 3D target-locking tracking

**DOI:** 10.1101/2024.09.25.614875

**Authors:** Hao Sha, Yu Wu, Yongbing Zhang, Xiaochen Feng, Haoyang Li, Zhong Wang, Xiufeng Zhang, Shangguo Hou

## Abstract

Fluorescence spectrum can provide rich physicochemical insights into molecular environments and interactions. However, imaging the dynamic fluorescence spectrum of rapidly moving biomolecules, along with their positional dynamics, remains a significant challenge. Here, we report a three-dimensional (3D) target-locking tracking-based single molecule fluorescence Spectrum Dynamics Imaging Microscopy (3D-SpecDIM), which is capable of simultaneously capturing both the rapid 3D positional dynamics and the physicochemical parameters changing dynamics of the biomolecules with enhanced spectral accuracy, high spectral acquisition speed, single-molecule sensitivity, and high 3D spatiotemporal localization precision. As a demonstration, 3D-SpecDIM is applied to real-time spectral imaging of the mitophagy process, showing its enhanced ratiometric fluorescence imaging capability. Additionally, 3D-SpecDIM is used to perform multi-resolution imaging, which provides valuable contextual information on the mitophagy process. Furthermore, we demonstrated the quantitative imaging capability of 3D-SpecDIM by imaging the cellular blebbing process. By continuously monitoring the physicochemical parameter dynamics of biomolecular environments through spectral information, coupled with 3D positional dynamics imaging, 3D-SpecDIM offers a versatile platform for concurrently acquiring multiparameter dynamics, providing comprehensive insights unattainable through conventional imaging techniques. 3D-SpecDIM represents a substantial advancement in single-molecule spectral dynamics imaging techniques.

## Main Text

Single-molecule fluorescence spectroscopy (SMFS) has become a prevalent tool for deciphering biomolecular interactions, conformation dynamics and molecular compositions^1–5^. The spatially or temporally separated molecular detection of SMFS enables measuring the non- equilibrium molecular dynamics that would otherwise be buried in ensemble average measurements. To date, the advancement of SMFS has paved the way for significant discoveries in various fields, including biochemistry, biophysics, and molecular biology. For instance, SMFS has elucidated the intricate dynamics of enzyme catalysis, protein folding, molecular tautomerization, and nucleic acid interactions with unprecedented precision^6–16^. In addition to leveraging fluorescence intensity information to probe the positional dynamics of biomolecules, the fluorescence spectrum provides rich physicochemical insights into the molecular environment and interactions^2, 17–26^. The spectral characteristics, including peak shifts, bandwidth, and emission profiles, reveal crucial details about the local microenvironment, binding events, and conformational states of the biomolecules under investigation. Interrogating the fluorescence spectrum dynamics can provide critical physicochemical parameter changing information of the studied biomolecules.

Recent years, various spectrally resolved single-molecule fluorescence imaging techniques have been developed to capture the distribution of biomolecular fluorescence spectra within cells^21, 22, 27–29^. To thoroughly investigate biological events, it is essential to continuously monitor the dynamics of the fluorescence spectrum, along with the biomolecules’ location changing dynamics. However, although advancements in spectral dynamics detection have been made^28, 30, 31^, imaging the fluorescence spectrum dynamics of rapidly moving biomolecules at high speed and with long observation time remains a significant challenge. This challenge arises primarily from multiple factors: (i) the rapidly moving target biomolecule may traverse outside of the excitation focal plane during the imaging period, which leads to insufficient spectrum dynamics data acquisition; (ii) the swift motion of target biomolecule during camera’s exposure time can result in the blurring of the fluorescence spectral image; and (iii) the limited number of fluorescence photons collected during short exposure times restricts the spatiotemporal localization precision of the biomolecules.

To overcome these limitations, we have developed a three-dimensional target-locking tracking-based single molecule fluorescence Spectrum Dynamics imaging Microscopy (3D- SpecDIM). In 3D-SpecDIM, the target biomolecule is maintained within the excitation volume via target-locking 3D single molecule tracking (TL-3D-SMT). This strategy effectively prevented the loss of rapidly moving target molecules during imaging. Varies TL-3D-SMT methods have been developed in recent years^32–46^. In here we utilized the 3D single molecule active real-time tracking (3D-SMART) method to perform target-locking 3D single molecule tracking, which has single-molecule sensitivity, high tracking speed, large imaging depth, and remarkable long tracking duration time^43^.

3D-SpecDIM can simultaneously capture the rapid 3D positional dynamics and fluorescence spectrum dynamics of biomolecules with enhanced spectral accuracy, high spectral acquisition speed, single-molecule sensitivity, and high 3D spatiotemporal localization precision. We demonstrate its capability by observing freely diffusing single fluorescent microspheres and single fluorescent dyes in solution. The spectral precision, temporal resolution and sensitivity of spectral imaging, and the spatiotemporal precision of 3D positional localization of 3D-SpecDIM have been thoroughly characterized. Furthermore, 3D-SpecDIM is applied to real-time spectral imaging of the mitophagy process, demonstrating its enhanced ratiometric fluorescence imaging capability. Additionally, 3D-SpecDIM enables multi-resolution imaging, providing valuable contextual information on the mitophagy process. Finally, 3D-SpecDIM has been employed to monitor the dynamics of cell membrane polarity changes during membrane blebbing, illustrating its capacity for multiparameter quantitative imaging.

## Results

### 3D-SpecDIM Overview

The general outline of 3D-SpecDIM is shown in **Fig. 1**. A target-locking 3D single molecule tracking (TL-3D-SMT) system is employed to maintain the target biomolecule within the excitation volume during data acquisition^43^. Within our TL-3D-SMT configuration, a pair of electro-optic deflectors and a tunable acoustic gradient index lens were utilized to drive the laser focus scanning in 3D (**Extended Data Fig. 1**). The fluorescence or scattering photons are collected by high-speed single-photon avalanche diodes and the photon arrival times were employed to compute deviation of the target molecule from the center of excitation volume in real-time on a field-programmable gate array. Subsequently, the feedback control voltages were applied to the piezo stages to relocate the molecule in the center of excitation volume and the 3D trajectory of molecule can be recorded. The TL-3D-SMT is able to capture the dynamics of single fluorophore with high speed and high spatiotemporal precision in 3D, which effectively prevented the loss of rapidly moving target molecules during data acquisition^41–43^. Additionally, the point-excitation- point-detection configuration of TL-3D-SMT facilitates observations at large imaging depths. To simultaneously acquire the single-molecule fluorescence spectrum, a prism-based spectral imaging system was integrated into the detection path^29, 47^. The fluorescence was dispersed via a prism and projected onto an electron multiplying charge-coupled device (EMCCD) to capture the spectral profile. The fluorescence was divided into reference channel and spectral channel through a beam splitter. The fluorescence in spectral channel was dispersed via a prism and projected onto one side of EMCCD and the fluorescence in reference channel was directly projected onto the other side of EMCCD. The pixel shifts between the images in the reference channel and spectral channel on EMCCD were utilized for registering the spectrum distribution. It is noteworthy that 3D- SpecDIM requires the capture of only one spectral stripe image at a time, enabling the use of a smaller EMCCD detection area and a higher frame rate. This capability significantly enhances high-speed spectral acquisition, achieving frame rate of up to 644 fps (**Supplementary Table 1**). After extracting spectral information from the spectral image, this data is synchronized with the 3D position of molecules with high temporal precision.

**Fig 1.**
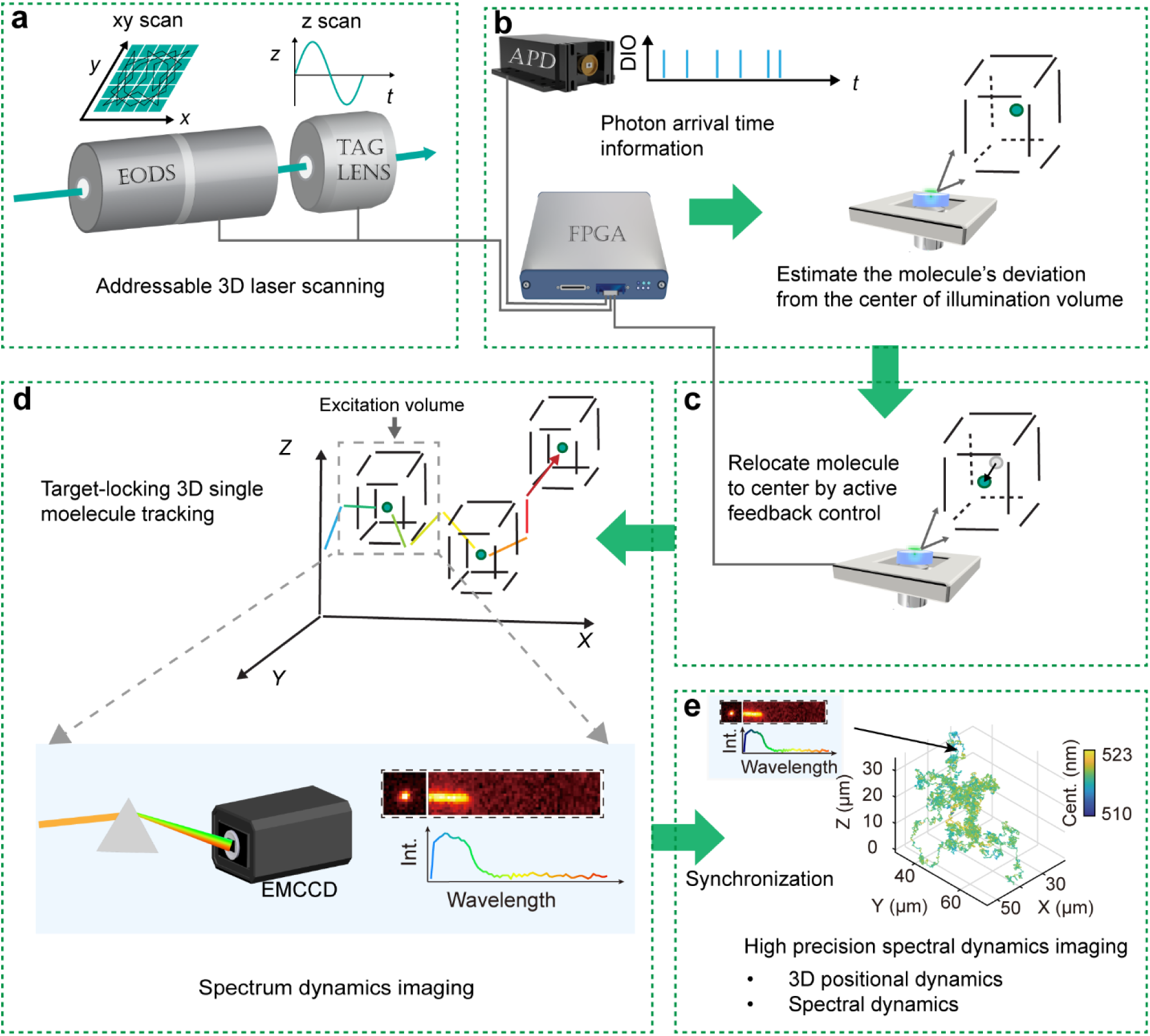
Schematic of 3D-SpecDIM. (**a**) A 2D EODs and a TAG lens are used to drive the focused laser spot rapidly scanning in a small volume (1 μm × 1 μm × 2 μm) after objective. (**b**) A FPGA utilizes the photon arrival time information collected by APD and the current laser position information to estimate the molecule’s deviation from the centre of illumination volume. (**c**) With the estimated molecule’s deviation information, a feedback control voltage is applied to the piezo stage to relocate the molecule to the centre of illumination volume. (**d**) During steps (**a-c**), the target molecule is continuously locked in the small excitation volume, resulting in high spatiotemporal precision 3D single molecule dynamics observation. Meanwhile, the target-locking imaging strategy allows for spectral acquisition with high imaging speed, large imaging depth and high spectral precision. (**e**) Synchronize the 3D positional dynamics and the spectral dynamics enables multiparameter dynamics acquisition.

### Characterization of the Performance of 3D-SpecDIM

The performance of 3D-SpecDIM was first characterized by spectrally tracking free diffusing 200 nm fluorescent microspheres in aqueous solution. The 3D positional dynamics of microspheres, along with their fluorescence spectrum dynamics can be simultaneously captured and synchronized (**Fig. 2a-b**). Moreover, we developed an optimized spectral detection method based on the Vision Transformer (ViT) model and domain adaptation strategy, which shows a significantly improvement on the spectral precision compared with the commonly used normal fitting method (**Fig. 2c, Extended Data Fig. 2** and **Extended Data Fig. 3**). It should be noted that the ViT-based spectral detection method performs more effectively in high signal situations than in low signal situations due to the influence of the signal-to-noise ratio (**Fig. 2d**). Subsequently, we characterized the spectral precision, spectral imaging speed and the sensitivity under various photon counts and exposure time situations with fixed microspheres (**Fig. 2d-f**). The results show that the spectral precision of 3D-SpecDIM can reach to 0.3 nm (**Fig. 2d**), while the temporal precision of spectral imaging can attain 1.55 ms (**Fig. 2e** and **Supplementary Table 1**). Moreover, the minimum emission rate required for successfully performing simultaneous 3D single molecule tracking and spectral imaging is 7 kHz (**Fig. 2f**), at the level of single fluorophore’s emission rate. Meanwhile, the spatiotemporal resolution of 3D positional localization can reach 1 ms and several nanometers (**Supplementary Fig. 1**). As shown in **Fig. 2d-f**, a trade-off exists between spectral precision and spectral imaging speed. By consulting these relationship plots, one can select appropriate imaging parameters based on the fluorescence emission rate of the sample and the specific requirements of the experiment.

**Fig. 2.**
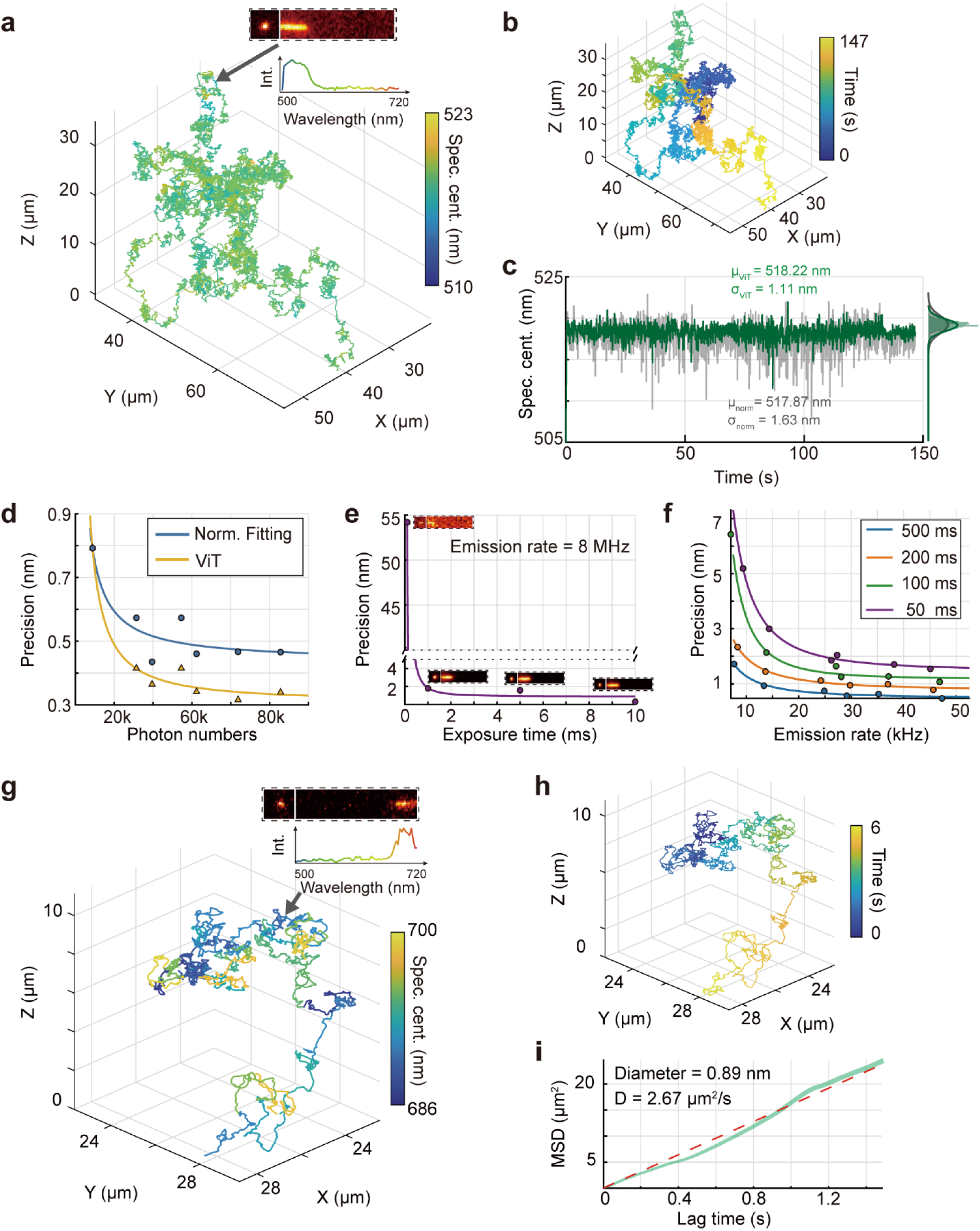
Characterization of the performance of 3D-SpecDIM. (**a**, **b**) 3D moving trajectory of a 200 nm fluorescent bead diffusing in water solution. The color indicates spectrum (**a**) or time (**b**). Insert in (**a**) shows the fluorescence spectral image in EMCCD and the spectral profile for one time point. (**c**) Comparison precision of normal fitting spectral localization method (gray line) and ViT spectral localization method (green line). Bin time: 29 ms. (**d**) Spectral tracking precision as a function of photon numbers. The blue line indicates the precision obtained with the normal fitting method, while the orange line shows the precision obtained with ViT method. Exposure time of EMCCD: 100 ms. (**e**) Spectral tracking precision as a function of exposure time. The emission rate of fluorescent bead is 8 MHz. (**f**) Spectral tracking precision as a function of emission rate with different exposure time. (**g, h**) 3D moving trajectory of a single SeTau 647 fluorescent molecule diffusing in 90 % glycerol. The color indicates spectrum (**g**) or time (**h**). Insert in (**g**) shows the fluorescence spectral image in EMCCD and the spectral profile for current time point. (**i**) Mean square displacement (MSD) as a function of lag time for trajectory (**g**). The red dash line shows linear fitting of MSD, producing a diffusion coefficient of 2.67 μm^2^/s. The calculated hydrodynamic diameter of molecule is 0.89 nm, which is consistent with the size of single fluorophore.

Next, the single-molecule spectrum dynamics imaging capability of 3D-SpecDIM was demonstrated by tracking single free diffusing fluorophore, SeTau 647, in 90% glycerol solution (**Fig. 2g-i**). Despite the low emission rate of the single fluorophore, we successfully conducted continuous and simultaneous monitoring of both its fluorescence spectrum and 3D positional dynamics for six seconds (**Fig. 2g-h**). The hydrated diameter of the fluorophore, calculated from the mean square displacement, was determined to be 0.9 nm, a dimension that falls within the single-molecule size range (**Fig. 2i**). This finding confirms the ability of 3D-SpecDIM to perform spectrum dynamics imaging at the single-molecule level. Notably, 3D-SpecDIM facilitates the acquisition of fluorescence spectra of freely diffusing molecules at picomolar concentration, representing a significant sensitivity enhancement over conventional fluorescence spectrometers. Additionally, this single-molecule spectral imaging approach also can significantly reduce the sample size required for analysis based on spectral change information.

### Identify particle switching with 3D-SpecDIM

In the realm of real-time single-particle tracking, a critical question frequently arises regarding the differentiation of two particles that exhibit similar fluorescence emission rates in one continuous tracking trajectory. We propose that this challenge can be addressed by analyzing differences in their fluorescence spectra. To validate this concept, we tracked mixed microspheres which have two distinct fluorescence spectra in solution using 3D-SpecDIM (**Extended Data Fig. 4** and **Supplementary Movie 1**). **Extended Data Fig. 4a** shows a trajectory where the tracked object transitioned from a yellow fluorescent bead to a green fluorescent bead. The two fluorescent beads exhibit similar fluorescent intensities, although the green bead displays greater intensity fluctuations. The switching event could not be distinguished from the trajectory alone, as there were no observable hopping movements (**Extended Data Fig. 4d**) and both particles have similar diffusion speed (**Extended Data Fig. 4c**). However, the particle switching can be easily identified from their spectral information. These results demonstrate that spectrum dynamics imaging can enhance the accuracy of real-time single particle tracking in complex environments.

### Improved ratiometric fluorescence imaging

Ratiometric fluorescence imaging is frequently used for monitoring the changes of local environment^48^. In traditional ratiometric fluorescence imaging, such as wavelength-split detection, two fluorescence channels are set for collecting fluorescence signals across different wavelength ranges. The ratio between the two channels is utilized as a means of spectral characterization. However, the accuracy of ratiometric imaging can be compromised by emission spectrum crosstalk, leading to reduced sensitivity and potential artifacts^49^. Fortunately, the capability of 3D- SpecDIM to acquire detailed spectrum profiles facilitates the implementation of spectral unmixing^50^, which significantly improves the sensitivity of ratiometric fluorescence imaging, as shown in simulation results (**Extended Data Fig. 5**).

To demonstrate the improved ratiometric fluorescence imaging capability of 3D-SpecDIM, we employed 3D-SpecDIM to monitor the mitophagy process (**Fig. 3a-g**). Mitophagy is an essential cellular mechanism that selectively degrades damaged or redundant mitochondria, thereby preserving cellular homeostasis and ensuring energy equilibrium^51^. Owing to the dynamic movement of autophagosomes and lysosomes during mitophagy, continuous observation of the mitophagy process within a singular focal plane presents significant challenges. The 3D target- locking tracking capability of 3D-specDIM effectively overcomes this by maintaining the autophagosomes in illumination volume, providing an unambiguously interaction dynamics information. During mitophagy, autophagosomes undergo translocation from the cytoplasm to the lysosome, wherein there is a decrease in pH (**Extended Data Fig. 6**). Therefore, we engineered a pH-sensitive fluorescent mitochondrial probe to monitor this translocation (**Fig. 3a, Supplementary Fig. 2, and Supplementary Fig. 3**). This probe integrates the mGold fluorescent protein, which exhibits a decrease in fluorescence intensity in response to diminishing pH levels, alongside HaloTag-JF549, whose fluorescence remains relatively stable across pH variations (**Supplementary Fig. 3**). Consequently, the intensity ratio of mGold to JF549, termed pH ratio, serves as an indicator of the local pH environment and was used to indicate the position of autophagosomes. Additionally, lysosomes are labeled with a deep red LysoTracker, allowing for monitoring the existence of lysosomes simultaneously.

**Fig. 3.**
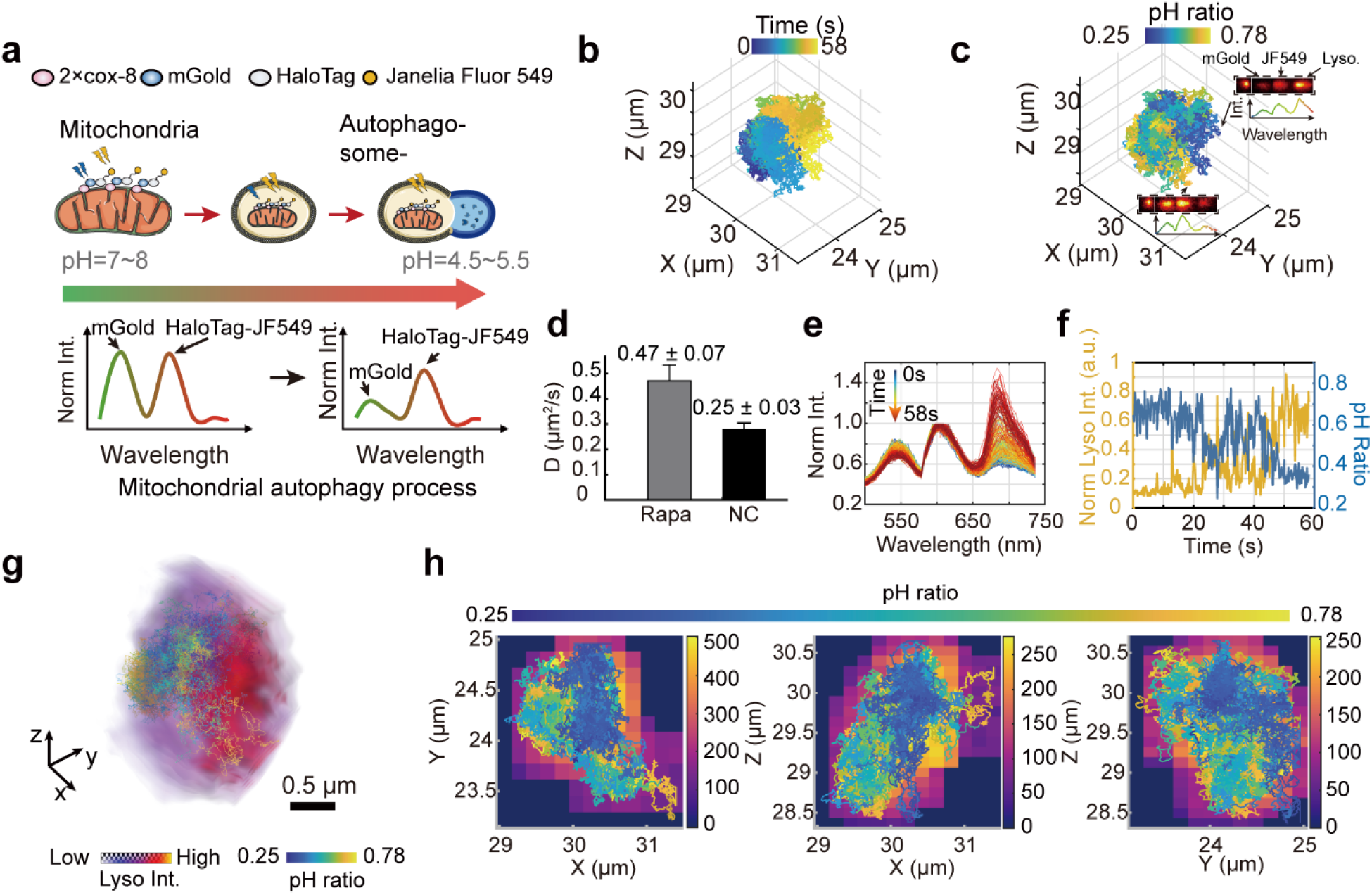
Enhanced ratiometric fluorescence imaging with 3D-SpecDIM in live cells. (**a**) Schematic of mitophagy. The mitochondrion was labeled with a pH-sensitive fluorescent probe mGold-HaloTag-JF549. The intensity ratio of mGold to JF549 serves as an indicator of the local environmental pH. As the pH of the local environment decreases, the intensity ratio correspondingly decreases. (**b, c**) 3D trajectory of a mitochondrion during mitophagy. The color indicates time (**b**) or intensity ratio (**c**). Inserts in (**c**) show the spectral images in two different time points indicated by arrows, which have different pH ratio values. (**d**) Comparison of mitochondrial moving speed in rapamycin-treated cell (0.47 ± 0.07 μm^2^/s) and untreated cell (0.25±0.03 μm^2^/s). n = 66. (**f**) The pH ratio of pH-sensitive fluorescent probe (blue line) or the intensity of lysosome (yellow line) as a function of time. Note that after 40 s, pH decreased when lysosome strength increased. (**e**) The fluorescence spectral profile change during mitophagy process. The three spectral peaks correspond to mGold, JF549, and LysoTracker, respectively. (**g**) Multi-resolution imaging of mitophagy process. The 3D volume image of lysosome is overlayed with spectrally encoded 3D moving trajectory of mitochondrion. The 3D lysosome volume image was reconstructed by registering the fluorescence photons to laser focus positions in a separated detection channel. Scale bar: 500 nm. (**h**) The sum intensity projection images of (**g)** in different planes.

We employed rapamycin (Rapa) to induce autophagy pathways via the inhibition of the mammalian target of rapamycin (mTOR). Subsequently, the autophagosome was tracked in 3D, while the spectra of mGold, JF549, and deep red LysoTracker were simultaneously monitored (**Fig. 3b-c**). The intensity ratio, derived from spectrum unmixing using 3D-SpecDIM, demonstrated a marked enhancement in both sensitivity and accuracy compared to wavelength- split detection methods, which rely directly on the intensity measurements from individual fluorescence channels (**Extended Data Fig. 5** and **Extended Data Fig. 7**). In rapamycin-treated cells, there was a notable increase in the number of trajectories exhibiting decreased pH ratio compared to the control group (17 versus 1 per hour), and these trajectories also show higher diffusion coefficients (**Fig. 3d-f** and **Supplementary Fig. 4**).

Additionally, 3D-SpecDIM has the capability to perform multi-resolution imaging. The fluorescence photons and the positions of the 3D scanning laser focus are recorded at each time point. By registering the fluorescence photons to laser focus positions in a separated detection channel, the 3D volume images of lysosome can be obtained while performing the spectral dynamics tracking. Multi-resolution imaging is achieved by aligning the 3D spectral trajectory with the 3D volume image, thereby providing valuable contextual information on the mitophagy process (**Fig. 3g-h** and **Supplementary Movie 2**).

### Multiparameter quantitative imaging

A distinctive feature of 3D-SpecDIM is its capacity to continuously monitor the dynamics of fluorescence spectral profile changes, while simultaneously acquiring 3D positional dynamics. This capacity facilitates the quantitative analysis of the kinetics and dynamics of biological interactions. Here we employed 3D-SpecDIM to perform quantitative imaging of the cellular blebbing process. Cellular blebbing, characterized by transient protrusions of the cell membrane, plays a crucial role in cell apoptosis and migration processes^52^. We observed that HeLa cells stained with Nile Red and exposed to high-intensity blue light exhibited a pronounced tendency to undergo membrane blebbing (**Supplementary Table 2**). To imaging this process, we employed 3D-SpecDIM to track the 3D position dynamics of 100 nm silver nanoparticles (AgNPs) attached to the cell surface^53^, simultaneously monitoring the fluorescence spectral dynamics of Nile Red. Nile Red is a solvatochromic, lipophilic fluorescent dye whose fluorescence emission spectra vary in response to the polarity of its environment. Therefore, it can serve as an indicator of cell membrane polarity^54^ (**Fig. 4a-c, Supplementary Fig. 5**, and **Extended Data Fig. 8** and 9). Notably, the blebbing frequently initiated at locations where AgNPs were present, potentially due to the plasmonic-enhanced electromagnetic field near the AgNPs surface^55^ (**Supplementary Movie 3**). The results show that the half accumulative probability time for bleb appearance is 5.26 minutes (**Fig. 4d**). Furthermore, blebbing occurred more frequently at the cell periphery than on the inner side, with a frequency 1.75 times higher (**Extended Data Fig. 9d**). Intriguingly, during blebbing events, AgNPs exhibited pronounced vertical moving speed and moving distance (0.45 ± 0.08 μm per minute), with less translational moving speed and moving distance (0.27 ± 0.02 μm per minute) (**Fig. 4e** and **Extended Data Fig. 10**), suggesting that the nanoparticles were elevated by the expanding bleb. During the cellular blebbing process, the membrane bleb size increased concomitant with a decrease in the spectrum peak position, indicating a decrease in membrane polarity, with a decreasing rate of 0.65 ± 0.32 nm per minute (**Fig. 4c**). In contrast, cells without blebbing showed considerably smaller changes in both the movement distance of AgNPs and the spectral dynamics **(Supplementary Table 3)**.

**Fig. 4.**
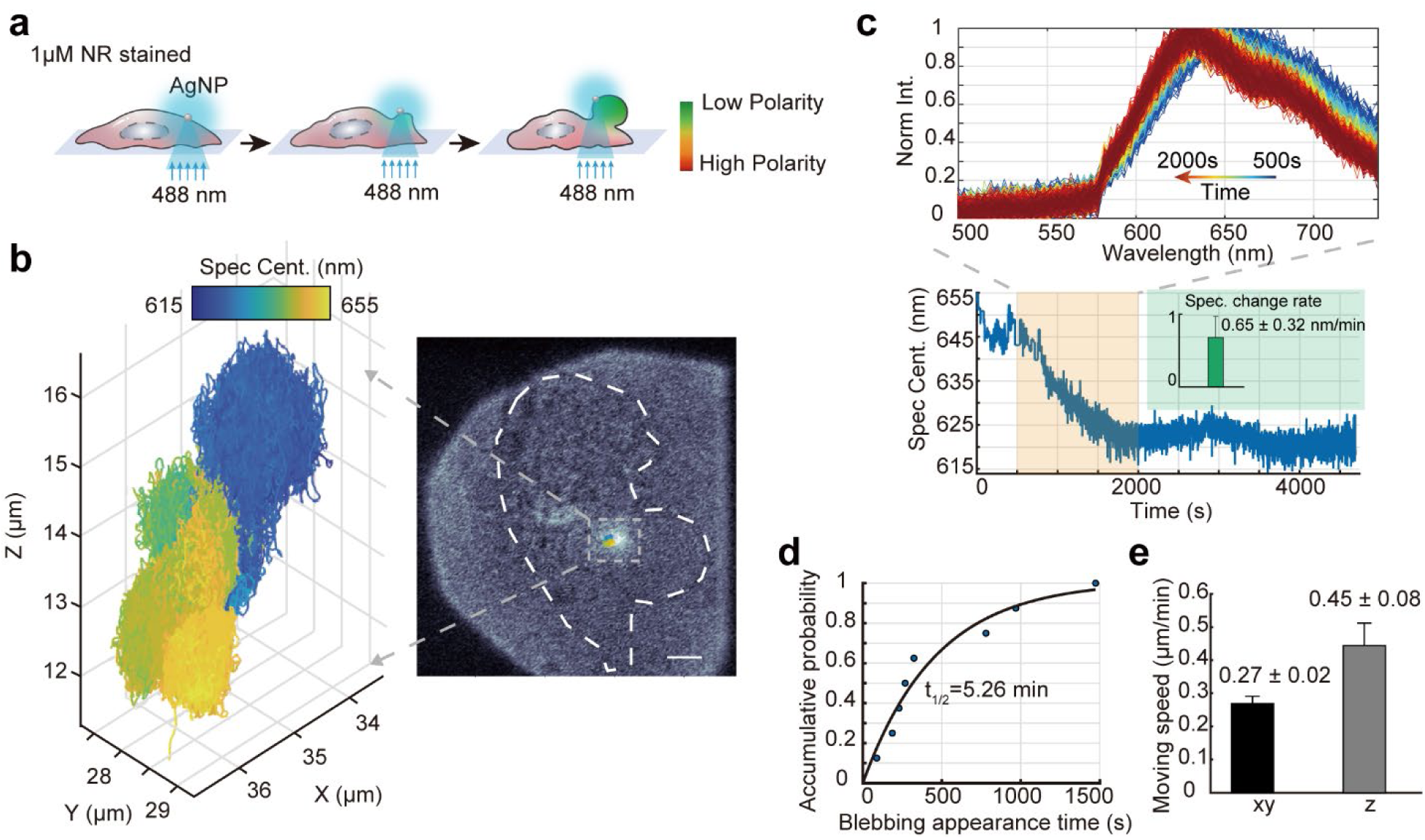
Quantitative spectral imaging with 3D-SpecDIM in live cells. **(a)** Schematic of cellular blebbing. The cell membrane was stained with Nile Red. With blue laser illumination, the cellular blebbing can be induced. AgNP was tracked in 3D and the fluorescence spectral dynamics of Nile Red was monitored by 3D-SpecDIM. (**b**) 3D trajectory of a single AgNP during cellular blebbing event. The color in trajectory indicates the fluorescence spectrum profile centroid position of Nile Red. Insert: The overlayed bright field image with 3D trajectory of AgNP. The white dash line shows the profile of cell. (**c**) Upper panel: the changing fluorescence spectral profiles of NileRed from 500 to 2000 seconds during the blebbing process; Lower panel: fluorescence spectral centroid position of Nile Red as a function of time during the cellular blebbing. Insert: The spectral change rate of Nile Red during cell blebbing. The spectral change rate is calculated for every 60 seconds with 8 trajectory data. n = 90. (**d**) The accumulative probability of cell blebbing appearance as a function of time. The points data were curve fitted with the function *y* = *a*(1 − *e*^−*b*⋅*x*^). When y = 0.5, the corresponding x value is 315.93s. (**e**) The comparison between vertical and translational moving speed of AgNPs during cell blebbing. n = 148.

## Conclusion

In summary, we developed 3D-SpecDIM, a real-time single molecule spectrum dynamics imaging method utilizing 3D target-locking tracking. This method enables simultaneously capturing the 3D location dynamics and fluorescence spectral dynamics of biomolecule with enhanced spectral accuracy, high spectral acquisition speed, single-molecule sensitivity, and high 3D spatiotemporal localization precision. The capability to detect multiple parameter dynamics allows for the interrogation of subtle physicochemical parameter changes within the microenvironment during rapid biological events. Furthermore, its capacity for spectral profile acquisition facilitates the use of spectrum unmixing to achieve higher sensitivity and accuracy in ratiometric fluorescence dynamics imaging. Additionally, the continuous detection of spectral changes, coupled with 3D positional dynamics imaging, significantly improves the quantitative analysis of both the kinetics and dynamics of biological systems. These advantages make 3D- SpecDIM particularly attractive as a unique technique for comprehensive analysis in both biochemical and biological systems.

While 3D-SpecDIM was demonstrated with fluorescence spectral dynamics, it is also applicable to scattering spectrum dynamics imaging. This capability renders it suitable for a broad range of single nanoparticle spectral analysis in an untethered state, such as monitoring catalytic and chemical reaction processes^56^. Furthermore, 3D-SpecDIM can also be extended to Raman imaging, allowing the simultaneous acquisition of 3D particle positional dynamics and Raman spectral dynamics^57^. Together, 3D-SpecDIM provides a versatile platform for the concurrent acquisition of multiple parameter dynamics, delivering comprehensive insights unattainable through conventional imaging techniques.

## Methods

### The optical setup of 3D-SpecDIM

The schematic diagram of 3D-SpecDIM is shown in Extended Fig. 1. Three lasers with wavelengths of 488 nm, 561 nm and 638 nm (LBX-488-60-CSB-PPA, LCX-561L-50-CSB-PPA and LBX-638-100-CSB-PPA, OXXIUS) were used for excitation. Three pairs of achromatic doublets lenses and three pinholes were utilized for spatial filtering for the three lasers. Two dichroic mirrors (T510lpxru and T588lpxr, Chroma) were used for combining laTsers. Then the laser polarization was tuned by a half waveplate (AHWP05M-580, Tholabs) and cleaned by Glan-Thompson polarizer (GLP10-A, Lbtek) before entering a pair of EODs (Model 310A, Conoptics Inc.). The laser beam was then expended by a pair of lenses (AC254-075-A-ML and AC254-300- A-ML, Thorlabs). A tunable acoustic gradient lens (TAGLENS-T1, Mitutoyo) was used to modulate the axial focus position. Then a pair of lenses (AC254-250-A-ML and AC254-150-A- ML, Thorlabs) were used to relay the laser to objective lens (HC PL APO 100x/1.40 OIL CS2, Leica). A dichroic mirror (ZT405/488/561/640 rpcv2, Chroma) was utilized for separating the excitation and emission light.

Subsequently, emission fluorescence was filtered by band-pass filter (refer to **Supplementary Table 4** for detailed filter configuration) and split by a beam splitter and directed into tracking and spectral imaging paths. In the tracking path, emission light was focused onto an APD using an achromatic lens (L11, AC254-050-A-ML, Thorlabs). In the spectral path, the fluorescence was further divided by a 30/70 splitter (BSS10R, Thorlabs). A CaF2 prism (PS863, Thorlabs) was used to disperse the fluorescent spectral signal. Finally, the beams were reflected by right-angle prism mirror (MRA25L-E02, Thorlabs) and focused with two achromatic lenses (L12, L13; AC254-150- A-ML, Thorlabs). An EMCCD (iXon Ultra 897, Andor) was used to record the spectral distribution. A two-axis piezo nanopositioner (Nano-PDQ275, Mad City Lab) and a z axis piezo nanopositioner (Nano-OPQ65, Mad City Lab) were used to move the sample and objective, respectively. A FPGA (PCIe-7858, National Instruments Corp.) was utilized to count the fluorescent signal, calculate the molecular position, control the nanopositioners and record the 3D trajectory of molecules. The synchronization of 3D single molecule tracking and spectral imaging was realized by instant triggering EMCCD with tracking signal or by aligning the start time point of tracking and spectral signal. The spectral registration method can be found in **Supplementary Fig. 6**. The laser power after the objective lens for each experiment can be found in **Supplementary Table 3**.

### Sample preparation and 3D-SpecDIM imaging

#### Fluorescent nanoparticle sample

We used two types of fluorescent spheres (FSDG002 and FSSY002, Bangs Lab) to calibrate the 3D-SpecDIM system. For fixed beads, we diluted the fluorescent bead stock solution into PBS at a ratio of 1:1000. For free moving beads, the bead stock solution was diluted into pure water at a ratio of 1:4000. The fluorescent bead samples were imaged or tracked according to the configuration in **Supplementary Table 3**.

#### Single SeTau 647 dye molecule sample

SeTau-647-NHS (K9-4149, SETA Biomedicals) was initially dissolved in DMSO at a concentration of 5 mg/ml, then aliquoted into 5 µL vials and stored at -80 °C. The dye was then diluted to 100 pM with distilled water. Subsequently, 1 mL of the dye solution was mixed with 9 g of glycerol (56-81-5, Aladdin) and added to glass-bottom culture dishes (801001, Nest) for 3D-SpecDIM experiments.

#### Mitophagy tracking

HeLa cells were cultured in DMEM (PM150223-500, Pricella) supplemented with 10% FBS (C04001-500, VivaCell) and 1% penicillin-streptomycin (PB180120, Procell), maintaining a 37 °C and 5% CO_2_ environment until reaching 60–80% confluency. Prior to transfection, cells were plated in glass-bottom dishes (801001, Nest) at densities promoting ∼70% coverage. The transfections were performed with Lipofectamine 3000 (L3000008, Invitrogen). After post-transfection for 14-18 hour, cells expressing 2×COX8-mGold-HaloTag were labelled with JF549 ligand (GA1110, Promega). Then immediately add rapamycin (HY- 10219, MedChemExpress) to sample and incubate for 2 hours. The COX8 gene (cloned from pCMV-mGold-Actin-C-18, MiaoLingBio P50209), representing Cytochrome C Oxidase Subunit 8, plays a crucial role in the respiratory chain by encoding cytochrome C oxidase and targeting a distinct leader peptide (MSVLTPLLLRGLTGSARRLPVPRAK)^58^. The mitochondrion was fused with mGold, a yellow fluorescent protein from pCMV-mGold-Actin-C-18 (MiaoLingBio; P50209), and Halotag (**Supplementary Fig. 2**). The cloning of 2×Cox8-mGold-HaloTag was synthesized by Tsingke Biotech (Beijing, China) by referencing mEmerald-Mito-7 (plasmid 54160, Addgene) using seamless cloning.

For three-color mGold-HaloTag-Lysosome tracking, cells were further incubated with 50 nM LysoTracker™ Deep Red (L12492, Thermo Fisher Scientific) working solution for 20 minutes at 37 °C to enable precise visualization. The subsequent co-localization imaging (**Supplementary Fig. 7** and **Supplementary Fig. 8**) or tracking of mGold, JF549, and LysoTracker Deep Red was performed via confocal fluorescence microscopy (LSM 980, Zeiss) or 3D-SpecDIM. The presented trajectory in **Fig. 3b-c** was part of a long trajectory with obvious positional jumping. The alignment procedure of the tracking trajectory on EMCCD image can be found in **Supplementary Fig. 9**.

### Spectral unmixing

To improve the sensitivity of ratiometric fluorescence imaging, the spectral unmixing was adopted by utilizing the spectrum profiles acquired with 3D-SpecDIM. The process involves solving the equation *M* = *S* ⋅ *C* + *E*, where *M* is the measured signal, *S* is the known fluorescence spectrum measured by fluorescence spectrophotometer, *C* is the coefficient matrix representing the contribution of each source, and *E* is the error. To solve for *C*, the least squares method is applied, resulting in *C* = (*S*^*T*^*S*)^−1^*S*^*T*^*M*.

### Quantitative imaging of the cellular blebbing

HeLa cells were seeded in glass bottom culture dishes (801001, Nest) and incubated for 18-20 hours at 37 °C in 5% CO2. For staining, cells were treated with 1 μM Nile Red in the culture medium and incubated for 25 minutes. Simultaneously, AgNPs (103716, 1 mg/mL, Xfnano) were prepared at a 20 µg/mL concentration in PBS, with ultrasonication for 5 minutes before adding them to cell sample. Following the Nile Red staining, cells were washed with cold PBS twice and cooled on ice for 5 minutes before adding the 20 µg/mL AgNPs solution to achieve a final in-cell concentration of 10 µg/mL. Then incubate cell on ice for 10 minutes. Finally, cells were washed with PBS and ready for imaging. During data collection, the brightfield light source was turned on for a short period to assess the cell’s blebbing status, which temporary interrupted the collection of fluorescence spectral data due to the poor signal-to-noise ratio. The spectral information during this period was filled with the spectral values of the previous time point. For all live cell imaging with 3D-SpecDIM, the sample temperature was maintained to 37 °C with a heating system (TC-1-100, Bioscience Tools).

### Data availability

The data that support the findings of this study are available from the corresponding author upon reasonable request.

## Acknowledgements

We would like to acknowledge support from the Shenzhen Medical Research Fund (B2301003), the National Natural Science Foundation of China (22204106, 62031023, 62331011), the Shenzhen Science and Technology Project (GXWD20220818170353009), the Evident & Shenzhen Bay Laboratory Joint Optical Microscopic Imaging Technology Development Program (S234602004-4), and the Guangdong Provincial Pearl River Talents Program (2021QN02Z631).

## Author Contributions

S. H. initiated the project. S. H., Z. Y. and H. S. conceived of the project. H. S. and Y. W. conducted the experiments. H. S., Y. W., Z. Y. and H. S. analyzed the data. X. F., H. L., and Z. W. contributed to build the microscope and data analysis. X. Z. contributed to sample preparation. S. H., H. S. Y. W. and Z. Y. wrote the manuscript. S. H. and Z. Y. supervised the project.

## Extended Figures

**Extended Data Fig. 1.**
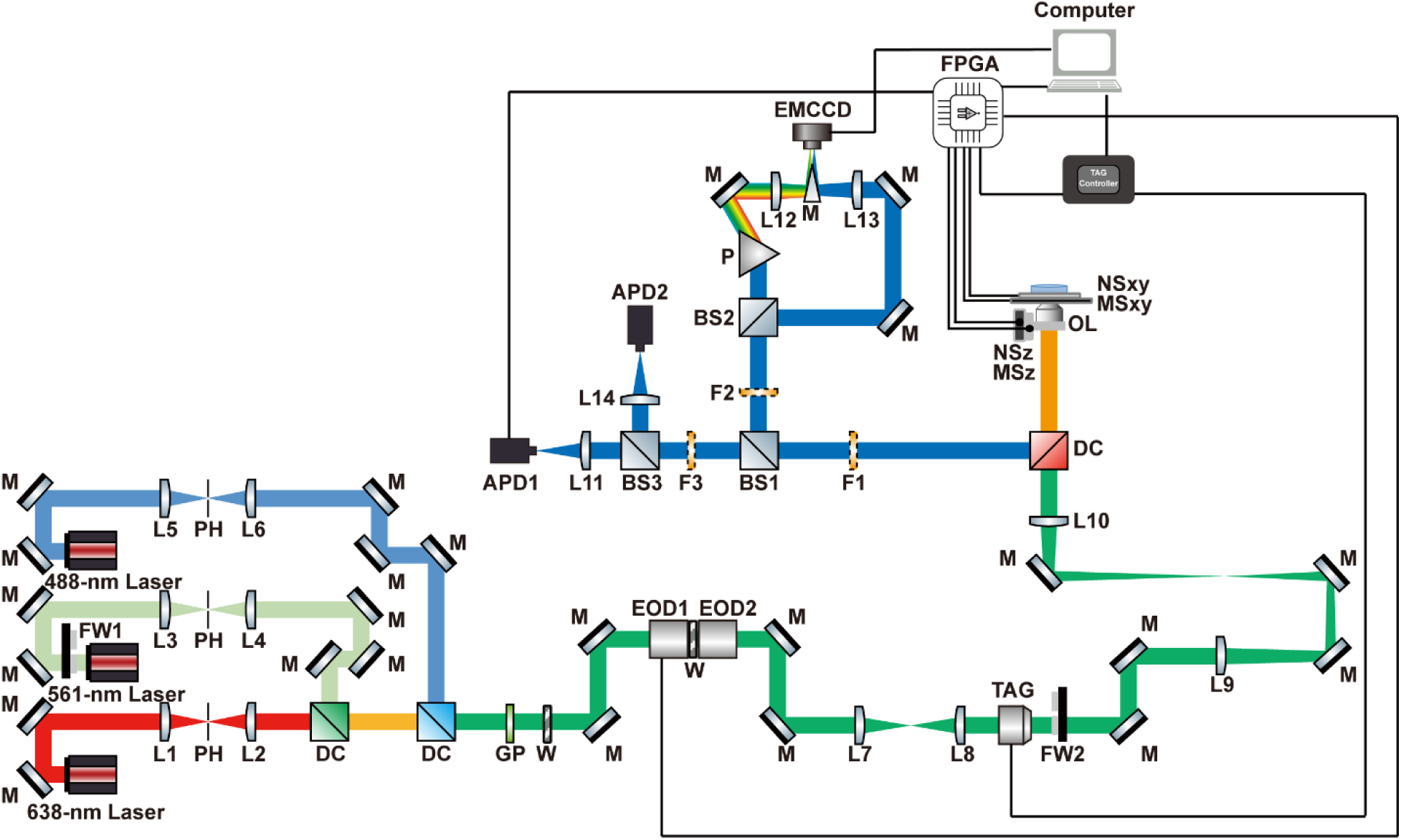
Schematic of 3D-SpecDIM setup. M: mirror; L: lens; PH: pinhole; FW: filter-wheel; PH: pinhole; DC: dichroic mirror; GP: Glan-Thompson polarizer; W: half waveplate; EOD: electro-optic deflector; TAG: TAG lens, OL: objective lens; MSxy: xy microstage; MSz: z microstage; NSxy: xy nanopositioner stage; NSz: z nanopositioner stage; FPGA: field programmable gate arrays; F: Fluorescence filter; BS: beam splitter; APD: avalanche diode detector; P: prism.

**Extended Data Fig. 2.**
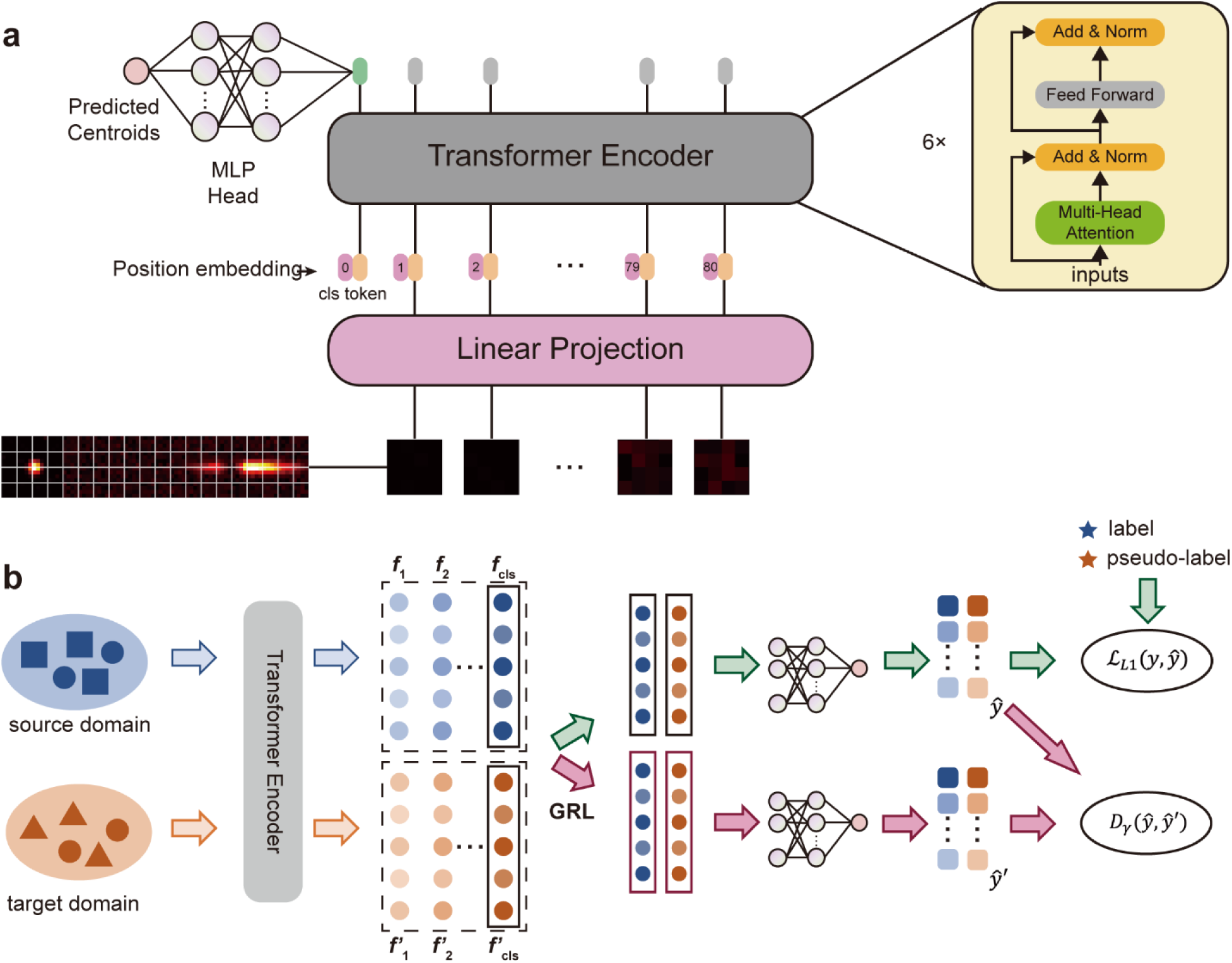
Vision Transformer and domain adaption-based spectral feature recognition. **(a)** Schematic of training process. Images with size of 16×80 are segmented into 4×20 patches using a 4×4 patch size, followed by the addition of learnable positional encodings after a linear projection layer. Several MLA (Multilayer Attention) layer iterations yield an output head, then spectral centroid features derived from a single-layer MLP (Multilayer Perceptron). **(b)** Inference process based on domain adaptive algorithm. For data that is unlabeled and falls outside the training data distribution, the system employs a gradient reversal layer (GRL) along with Margin Disparity Discrepancy (*D*_*γ*_(*ŷ*,*ŷ*′)) loss to finely tune the parameters of the Encoder. See **Supplementary Note 2** for detailed information.

**Extended Data Fig. 3.**
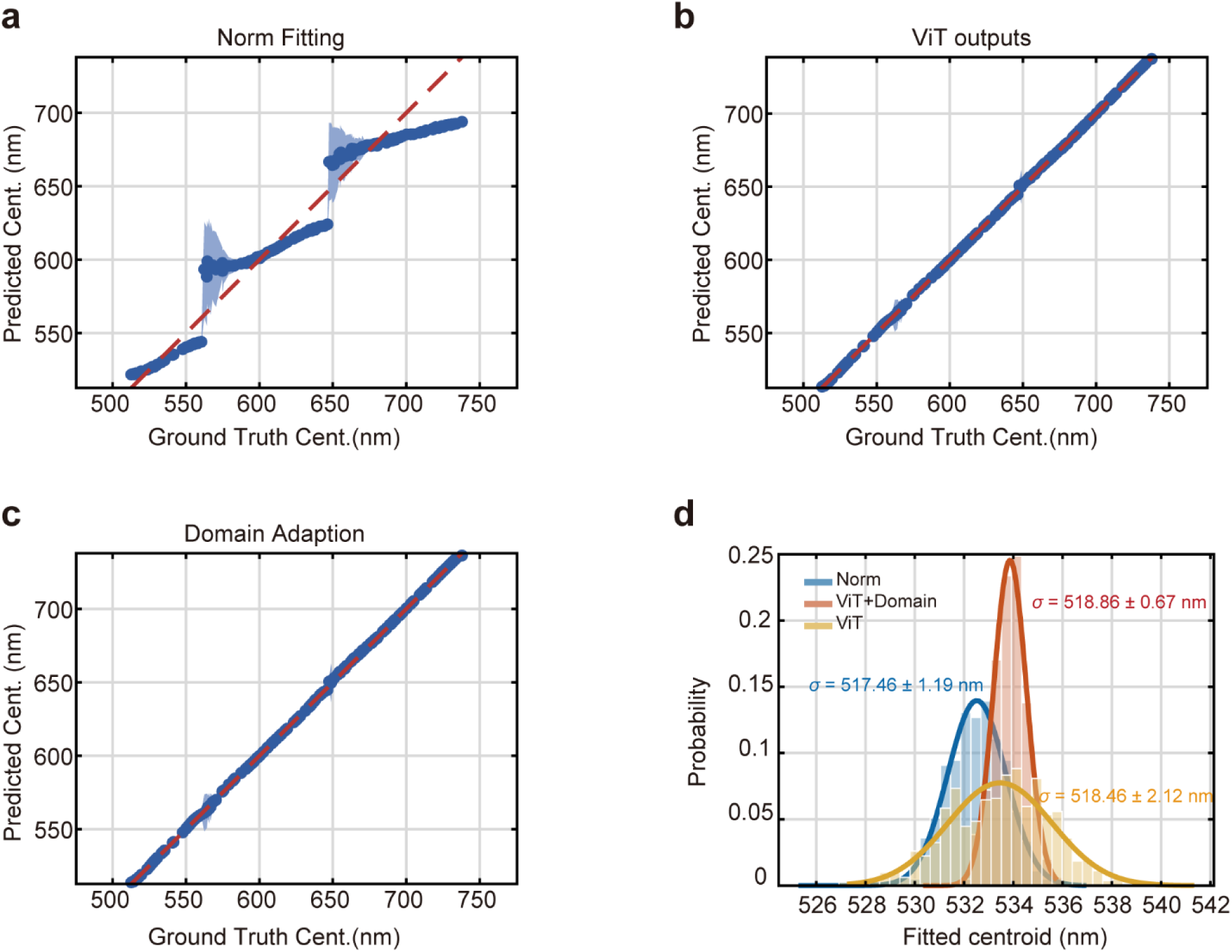
Comparison of centroid recognized methods. (**a-c**) Comparison the spectral localization precisions of **(a)** traditional normal fitting method, **(b)** Vision Transformer without domain adaption strategy, and **(c)** Vision Transformer with domain adaption strategy on simulation data sets. **(d)** Comparison the spectral localization precisions of different methods on fluorescent bead spectra data. The Vision Transformer with domain adaption strategy shows improved spectral precisions.

**Extended Data Fig. 4.**
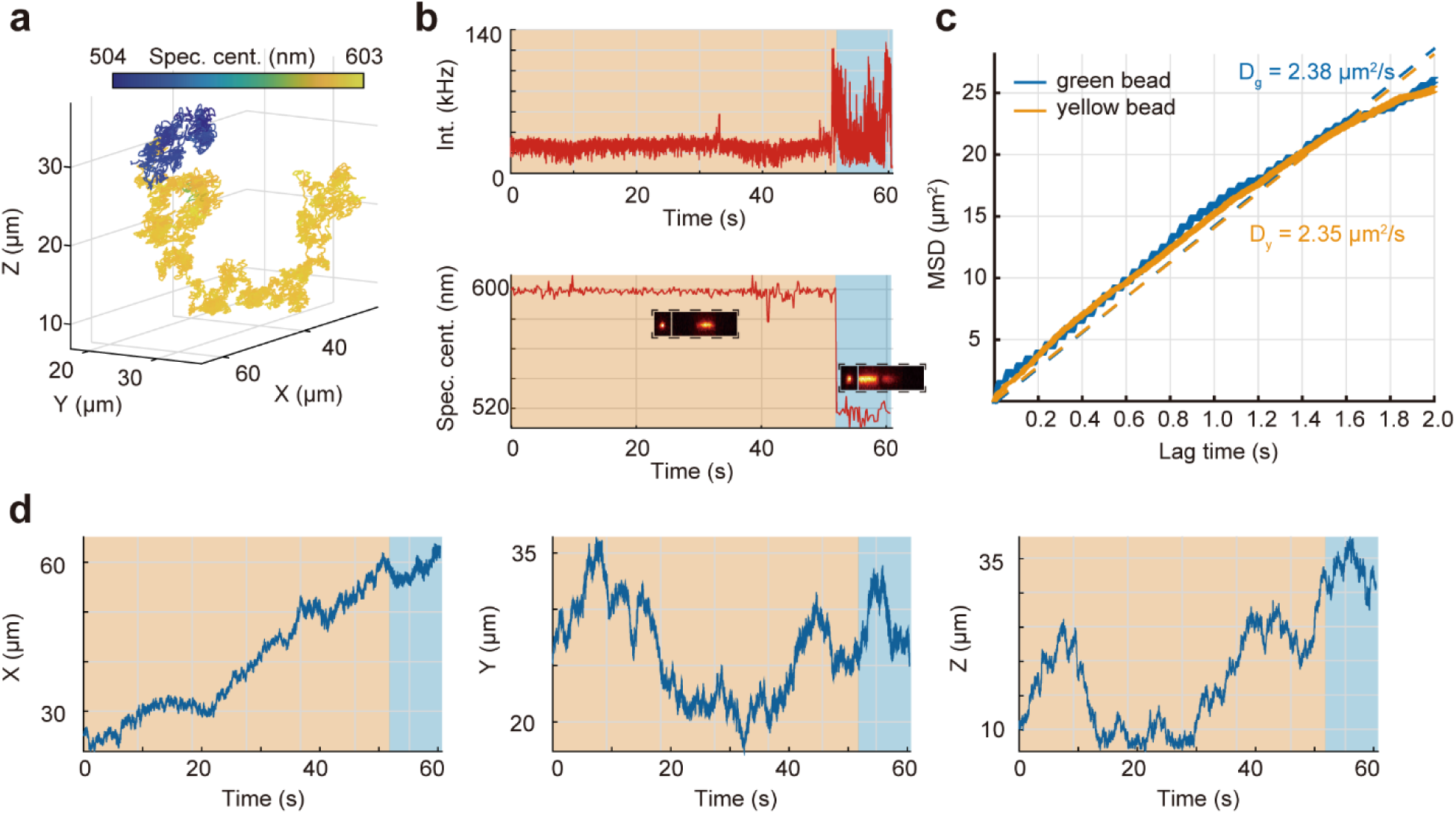
Discern two particles switching events during real-time single-particle tracking with 3D-SpecDIM. Green and yellow fluorescent beads were mixed in solution and tracked with 3D-SpecDIM. (**a**) 3D trajectory of a particle switching event. The trajectory of yellow fluorescent particle and green fluorescent particle are encoded with yellow color and blue color, respectively. (**b**) The fluorescence intensity (upper panel) and the fluorescence spectral centroid (lower panel) as a function of time in trajectory (**a**). Inserts show the spectrum image in EMCCD. (**c**) The mean square displacements as a function of time for yellow particle and green particle. The diffusion coefficients of them are 2.35 µm^2^/s and 2.38 µm^2^/, respectively. (**d**) The x, y, and z position as a function of time in trajectory (a).

**Extended Data Fig. 5.**
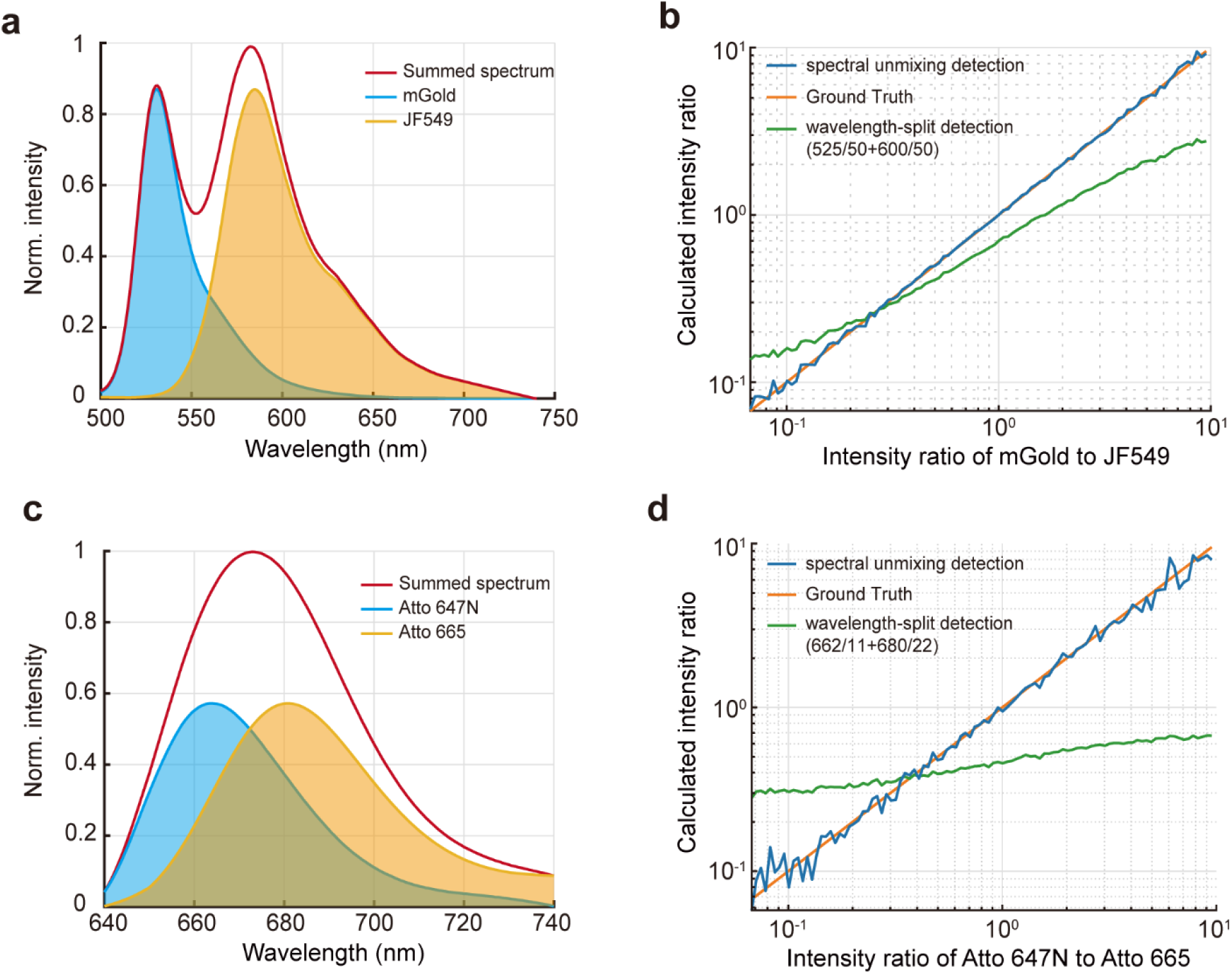
Spectral profile acquisition of 3D-SpecDIM enables high precision ratiometric fluorescence imaging by spectral unmixing. **(a)** The fluorescence emission spectrum of mGold and JF549 dye. The red line is the summed spectral intensity profile with both dyes have the peak intensity of 1. **(b)** Simulated comparison of wavelength-split detection method and 3D-SpecDIM enabled spectral unmixing in ratiometric fluorescence imaging. In wavelength-split detection method, two band pass filters (525/50 and 600/50) were adopted to collect the fluorescence of mGold and JF549 dye. The intensity ratio is calculated with the sum of fluorescence signal within the wavelength range defined by the band pass filters. In spectral unmixing method, fluorescence spectral profiles of mGold and JF549 dye were recovered from the summed spectral profile. The intensity ratio is calculated as the ratio of their spectral profile peak values. We set a various of intensity ratio of mGold to JF549 and compared the wavelength-split detection method (green line) and spectral unmixing method (blue line). The result shows that the 3D-SpecDIM enabled spectral unmixing method demonstrated pronounced precision enhancement. (**c**, **d**) Similar analysis with (**a**, **b**) but using a larger spectrum-overlapping fluorescent dye pair, Atto 647N and Atto 665.

**Extended Data Fig. 6.**
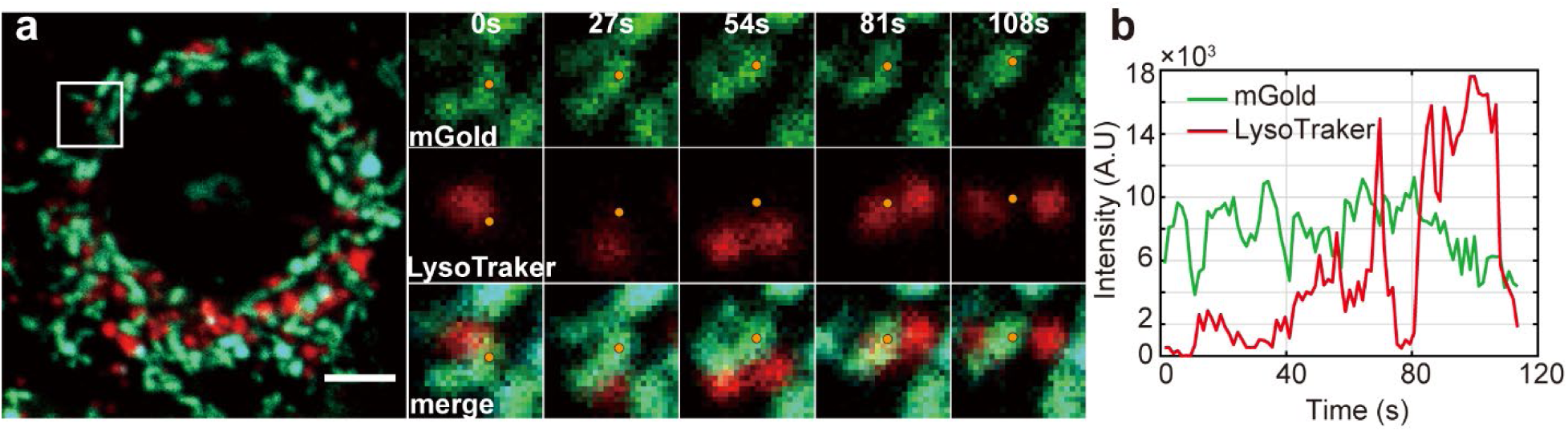
Monitor the mitophagy process with confocal microscopy. (**a**) Time-lapse confocal imaging of mitochondria and lysosome during mitophagy process, with mitochondria show in green and lysosomes show in red. Scale bar = 5 µm. **(b)** The fluorescence intensity of mGold on mitochondria and lysosome as a function of time in positions marked with orange dot in (a). The mitochondria were labeled with mGold and the lysosomes were labeled with LysoTracker Deep Red. The orange dot positions were determined by finding the largest fluorescence intensity pixel in each mitochondrion image. The intensity of mGold was decreased when mitochondria merged with lysosome.

**Extended Data Fig. 7.**
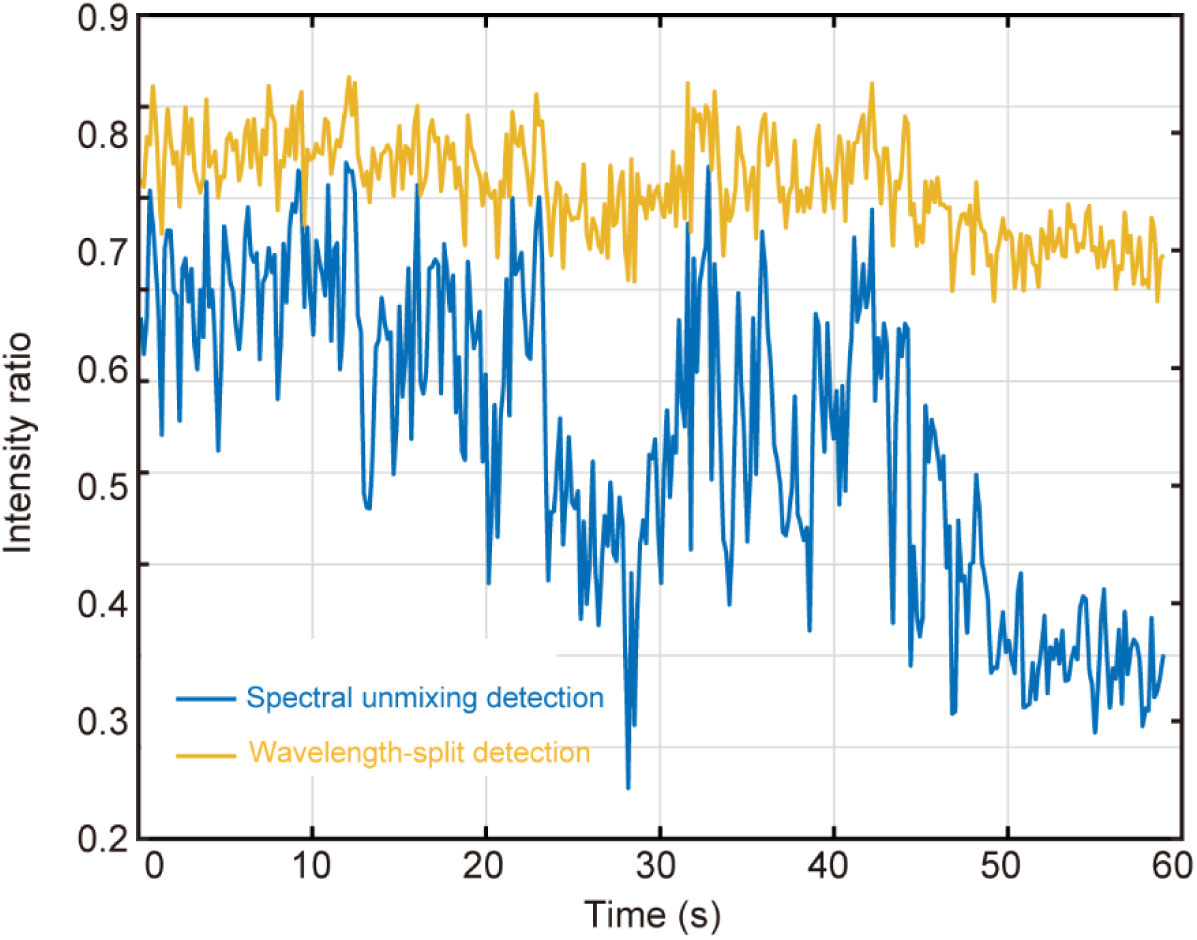
Comparison of the ratiometric fluorescence detection between wavelength-split detection (yellow line) and spectral unmixing detection (blue line) methods. The spectral unmixing detection method shows much larger changing range, demonstrating improved sensitivity.

**Extended Data Fig. 8.**
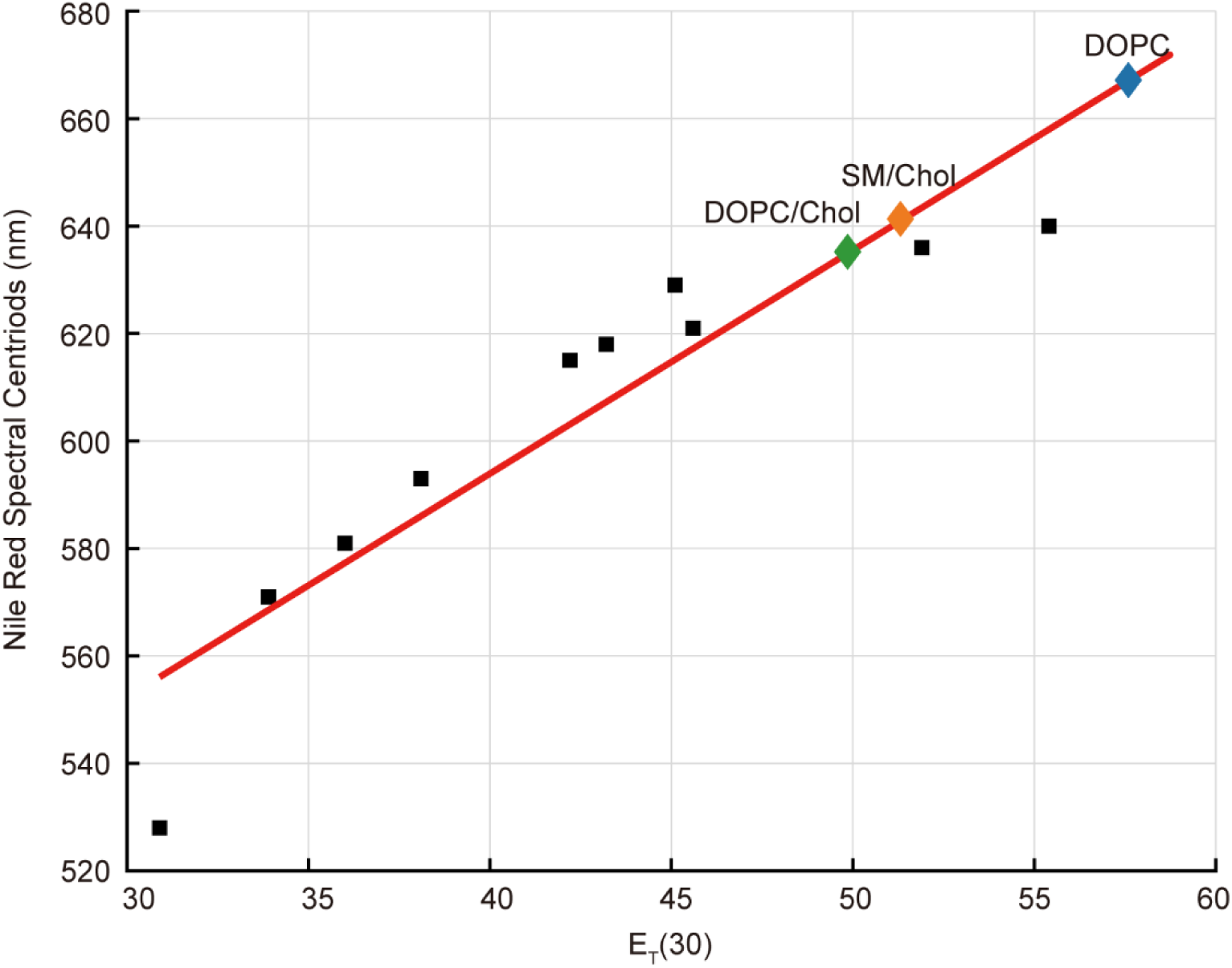
The relationship between spectral centroid of Nile Red and E_T_(30), a commonly used parameter for quantifying the polarity. The black square dots indicate the Nile Red spectral centroid position in different solution with corresponding E_T_(30), a parameter used for characterizing the polarity of solvent. The green diamond dot, orange diamond dot and blue diamond dot show the Nile Red spectral centroid position in different solutions as marked on the plot figure. Therefore, the polarity of these solution, or the value of their E_T_(30), can be extracted.

**Extended Data Fig. 9.**
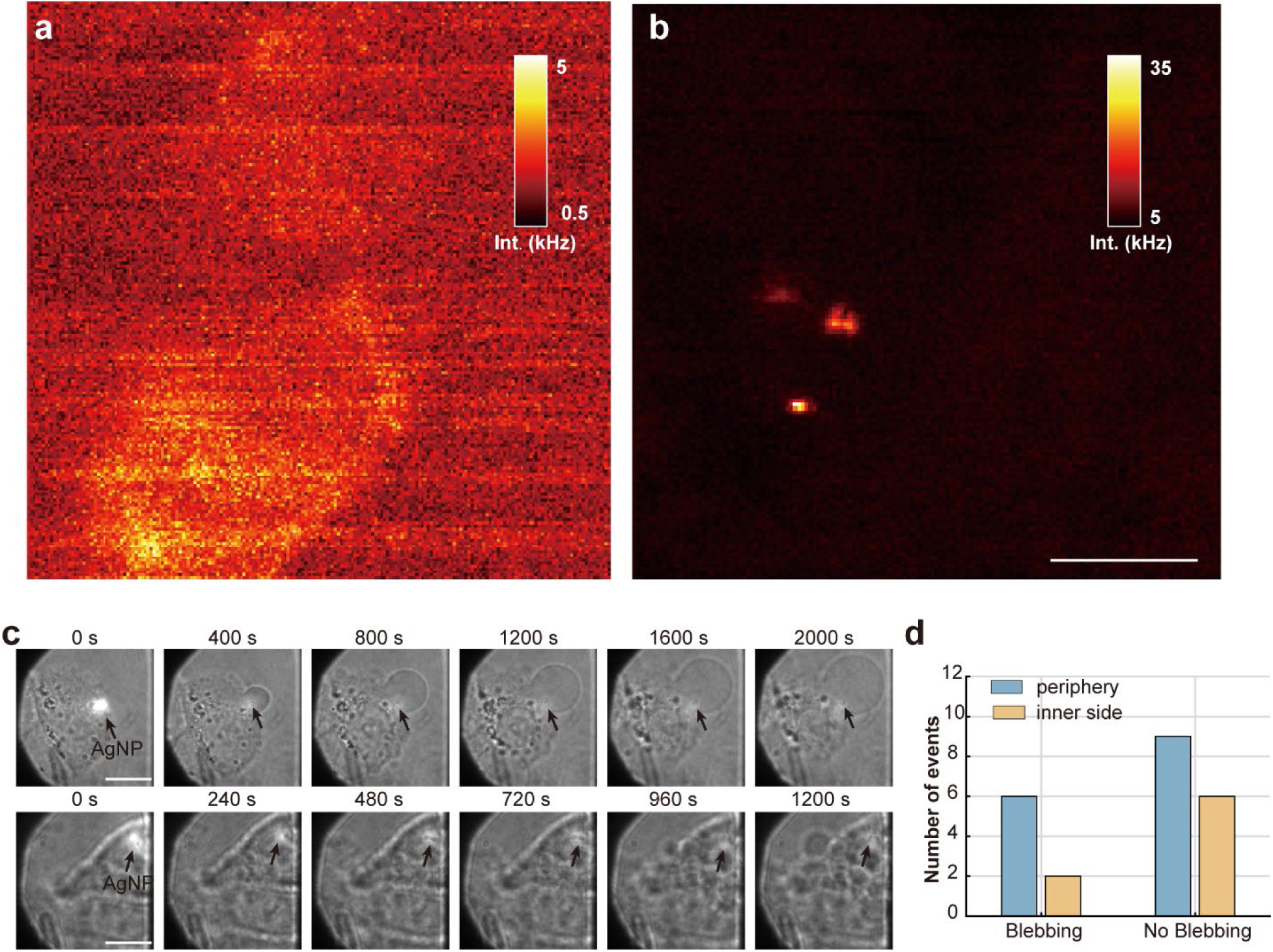
Signal to background ratio characterization of AgNPs scattering imaging in cell and the bright field images of cellular blebbing process. **(a)** Scanning image of Hela cells without AgNPs. Hela cells were labeled with Nile Red. The 488 nm laser was used for excitation and a 488/10 nm bandpass filter coupled with a polarizer were used in the detection path. **(b)** Scanning image of Hela cells with AgNPs. **(c)** Time-lapse bright field images of cellular blebbing imaging process. The upper panel shows cellular blebbing occurred at the periphery of cell, while the lower panel shows cellular blebbing occurred on the inner side of cell. Scale bar: 10 μm. (**d**) Blebbing occurred frequency at different cell position (n = 24). We examined the blebbing positions occurred on cell. The results show that blebbing occurred more frequently at the cell periphery (43.75%) than on the inner side (25%).

**Extended Data Fig. 10.**
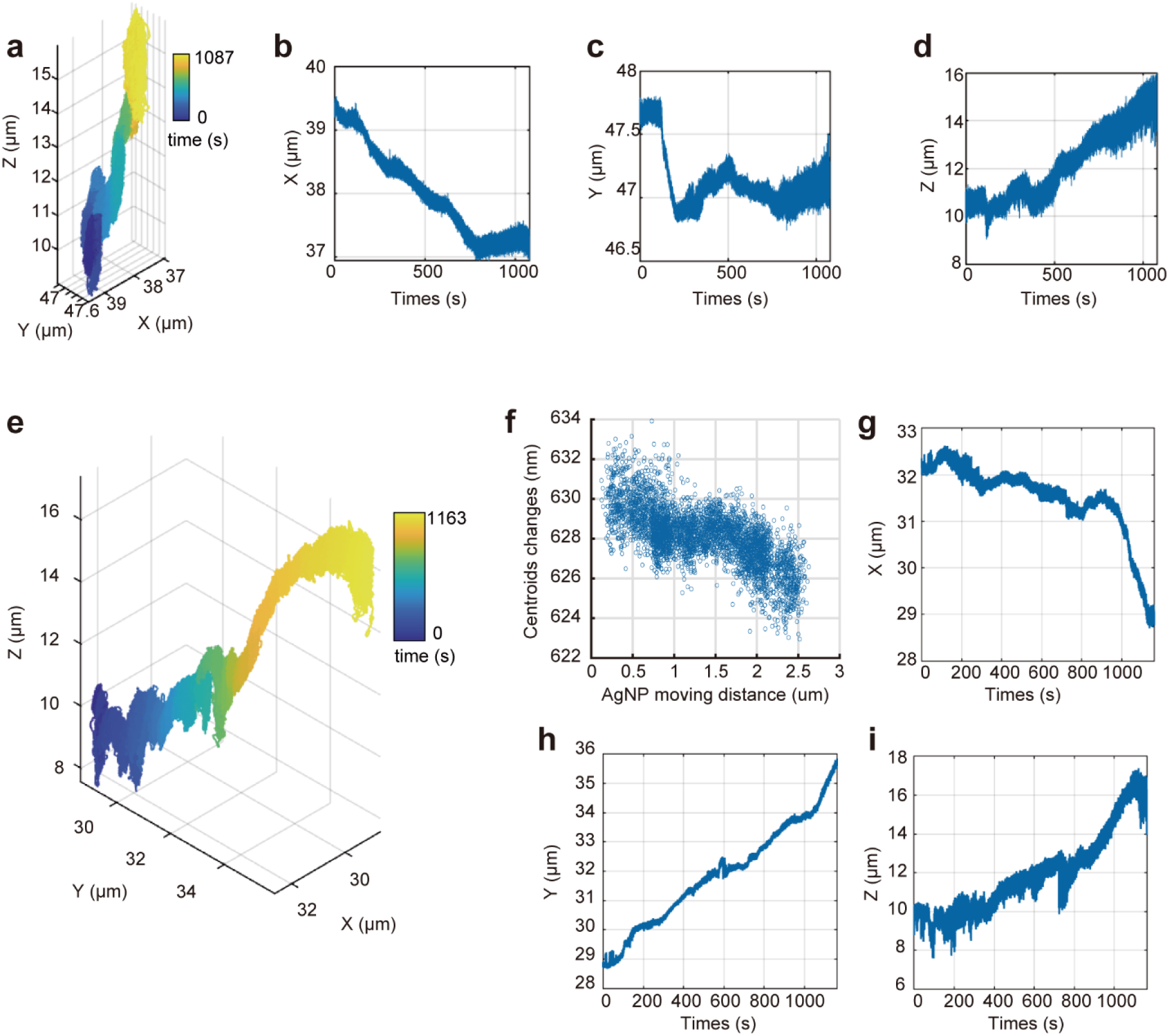
3D trajectories of AgNPs during celluar blebbing. **(a)** 3D trajectory of AgNP during blebbing. The AgNP initially located on the inner side of cell. (**b-d**) x, y, and z position of AgNP as a function of time. **(e)** 3D trajectory of AgNP during blebbing. The AgNP initially located at the periphery of cell. (**f**) Fluoruescnece spectrum centroids changes of Nile Red as a function of AgNP moving distance. (**f**-**h**) x, y, and z position of AgNP as a function of time.

## Movie Description

**Supplementary Movie 1**. **Discern two particles switching events during real-time single-particle tracking with 3D-SpecDIM.** The trajectory of yellow fluorescent particle and green fluorescent particle are coded with yellow color and blue color, respectively. The particle swithing can be observed at approximately 51 seconds. The video is played at 5.5× speed.

**Supplementary Movie 2**. **Multi-resolution imaging of mitophagy process**. The 3D lysosome volume image is overlayed with spectrally encoded 3D moving trajectory of mitochondrion. The trajectory’s color represents the pH ratio. The right panel shows the real-time spectral images of the mGold-HaloTag-JF549 probe. Lysosome intensity was reconstructed using Avizo software and displayed using temperature-based color coding. The video is played at 6.4× speed.

**Supplementary Movie 3**. **Time-lapse bright field movie of cellular blebbing imaging process**. The cellular blebbing occurred at the periphery of the cell. The white arrow points to the tracked AgNP particle. The video is played at 52.5× speed.

