## Supplementary information for "Single molecule spectrum dynamics imaging with 3D target-locking tracking"

**Sha et al.**

| Section | Page |
|---|---|
| **Figures** | |
| Supplementary Figure 1. 3D tracking localization precision. | 1 |
| Supplementary Figure 2. Domain structure of 2×COX8-mGold-HaloTag used in mitophagy experiments. | 2 |
| Supplementary Figure 3. pH titration of mGold and JF549 dye. | 3 |
| Supplementary Figure 4. Spectrum dynamics tracking of pH-sensitive probe labeled mitochondrion in cells without mitophagy. | 4 |
| Supplementary Figure 5. Characterize the polarity of lipids membrane with 3D-SpecDIM. | 5 |
| Supplementary Figure 6. Spectral registration. | 6 |
| Supplementary Figure 7. Confocal images of mitochondria and lysosome during mitophagy. | 7 |
| Supplementary Figure 8. Comparison between 2×cox8-mGold-HaloTag and mitochondrial dyes. | 8 |
| Supplementary Figure 9. Align the tracking trajectory position with EMCCD image. | 9 |
| Supplementary Figure 10. Simulated datasets for ViT model training. | 10 |
| **Tables** | |

| | |
|---|---|
| Supplementary Table 1. Temporal resolution analysis of 3D-SpecDIM system. | 11 |
| Supplementary Table 2. Cell blebbing occurrence analysis under different conditions. | 12 |
| Supplementray Tables 3. Statistics data of fluorescence spectrum centroid changes during cell blebbing. | 13 |
| Supplementary Table 4. Optical path configurations and experimental parameters | 14 |
| Supplementary Table 5. Preparation of Silicon spheres coated with SLB. | 16 |
| **Notes** | |
| Supplementary Note 1. Time resolution analysis of 3D-SpecDIM system. | 17 |
| Supplementary Note 2. Cell blebbing occurrence analysis under different conditions. | 19 |
| Supplementary Notes 3. Multi-resolution imaging with 3D-SpecDIM. | 24 |

### Supplementary Figures

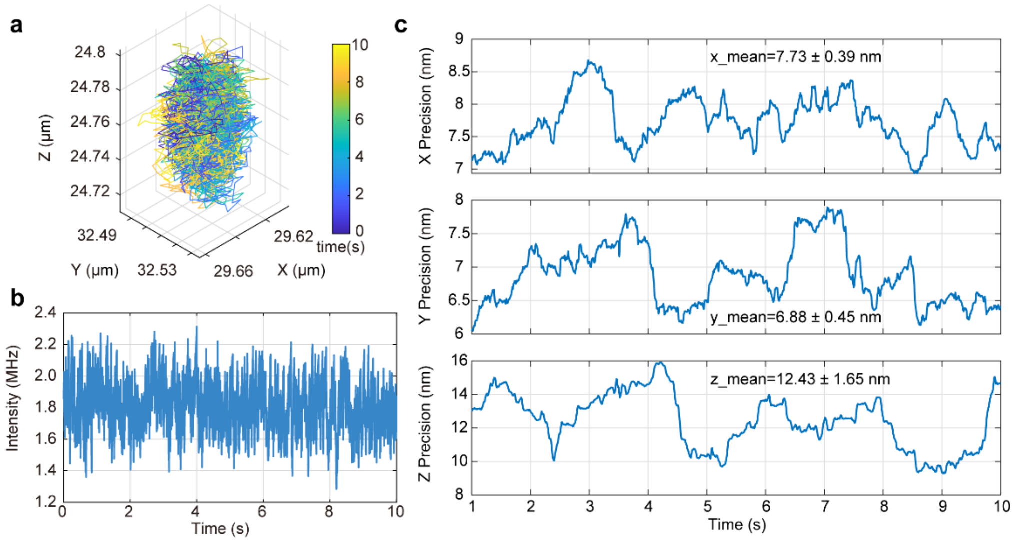

**Supplementary Figure 1. 3D tracking localization precision.** (**a**) 3D trajectory of a fixed 200 nm fluorescence bead. (**b**) Fluorescence intensity as a function of time. (**c**) Tracking precision as a function of time for X, Y, and Z, respectively. The precision is measured by the standard deviation of position of 1 s data with 10 ms sliding window.

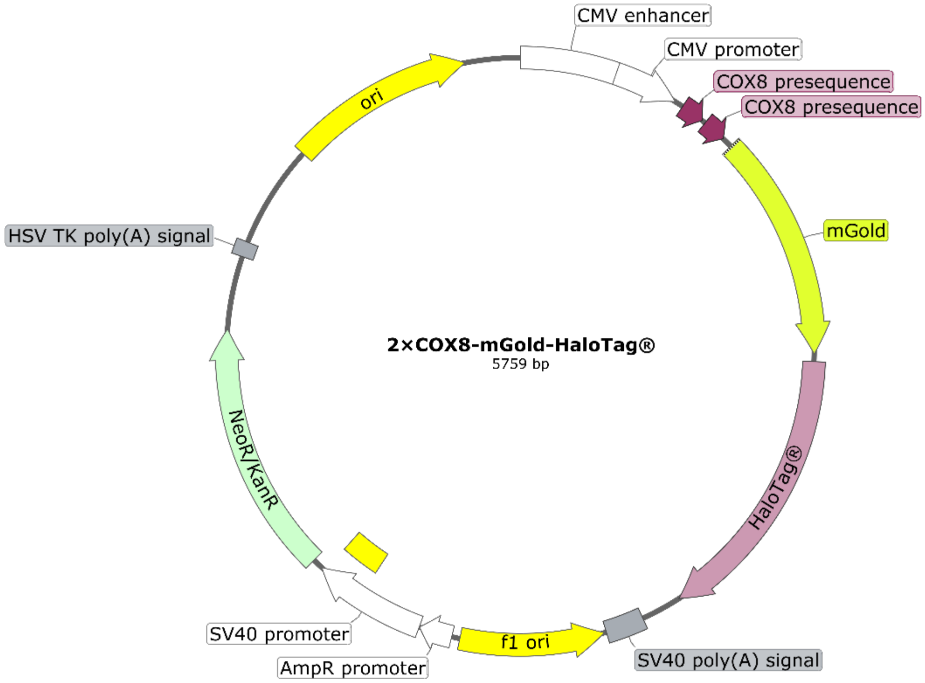

**Supplementary Figure 2. Domain structure of 2×COX8-mGold-HaloTag used in mitophagy experiments.**

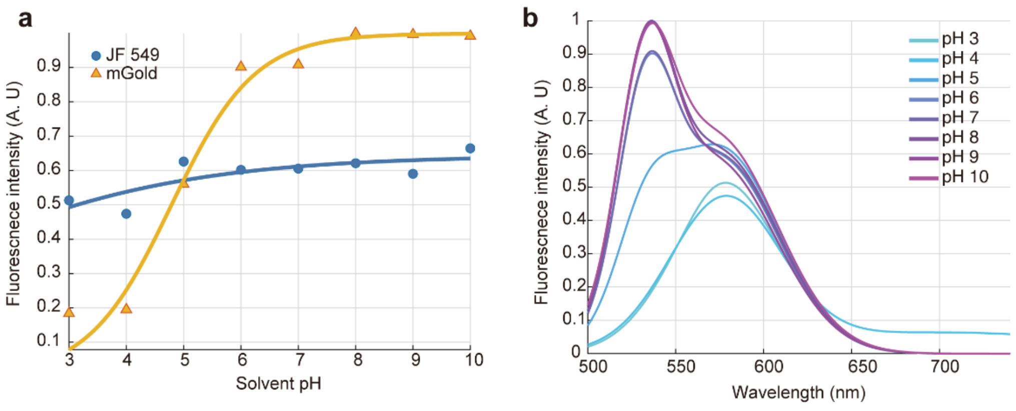

**Supplementary Figure 3. pH titration of mGold and JF549 dye.** (**a**) The fluorescence intensity of mGold (yellow triangle) and JF 549 (blue dot) as a function of pH of solvent. (b) The fluorescence spectral profile of mGold-JF549 in solutions with various pHs. Due to the differences in the absorption spectra of mGold and JF549, the combined spectrum in various pH buffers[1] was synthesized under 488 nm and 540 nm laser excitation, respectively. Specifically, after measuring the spectra of the mGold-JF549 mixture under 488 nm and 540 nm laser excitation separately, the spectra shown in (**b**) were obtained by summation and normalization.

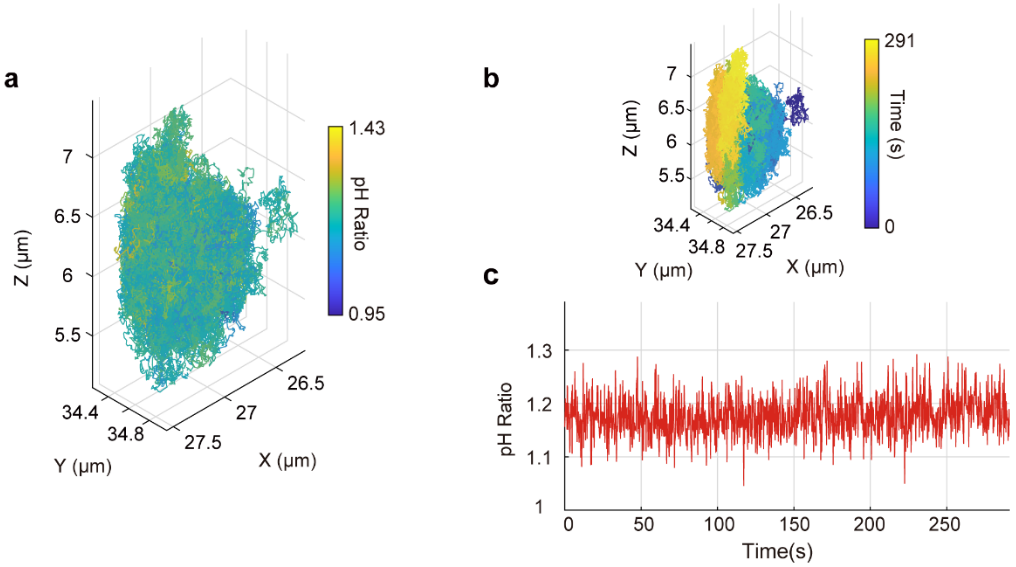

**Supplementary Figure 4. Spectrum dynamics tracking of pH-sensitive probe labeled mitochondrion in cells without mitophagy**. (**a**, **b**) 3D trajectory of a mitochondrion in live cell. The color indicates pH ratio (**a**) or time (**b**). (**c**) pH ratio as a function of time. The excitation laser power of 488 nm, 561 nm, and 638 nm are the same with experiment in **Figure 3f** in main text.

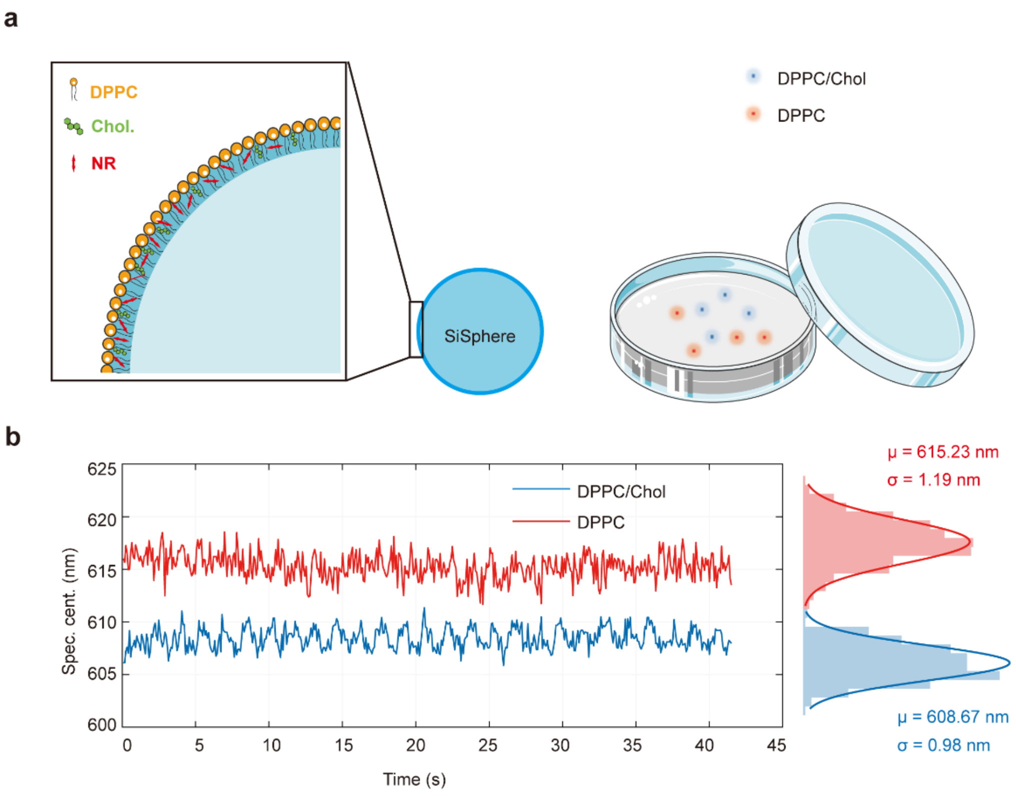

**Supplementary Figure 5**. **Characterize the polarity of lipids membrane with 3D-SpecDIM**. (**a**) Schematic diagram of the structure of SLB-coated 100 nm silicon spheres, where DPPC represents 1,2-dipalmitoyl-sn-glycero-3-phosphocholine, Chol represents cholesterol, and NR represents Nile Red. (b) With the existence of cholesterol, the polarity of lipids membrane was decreased, and the fluorescence spectrum shows blue shift.

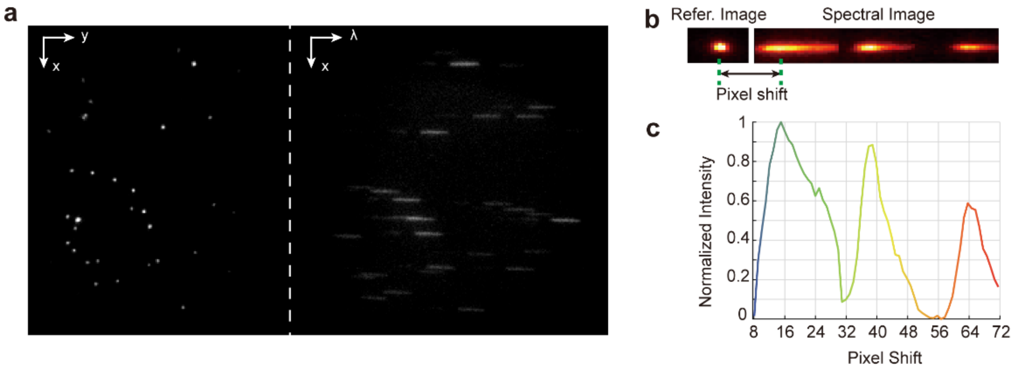

**Supplementary Figure 6. Spectral registration. (a)** The reference image (left) and spectral image of 200 nm yellow fluorescent microspheres on EMCCD. (**b**) The reference image (left) and spectral image (right) of four-colors fluorescent bead (Thermo Fisher Scientific, TetraSpeck™, T7279) on EMCCD. The spectral image was generated by horizontally flipping the raw spectral image. (**c**) Fluorescence intensity distribution versus pixel shift for spectral image (**b**).

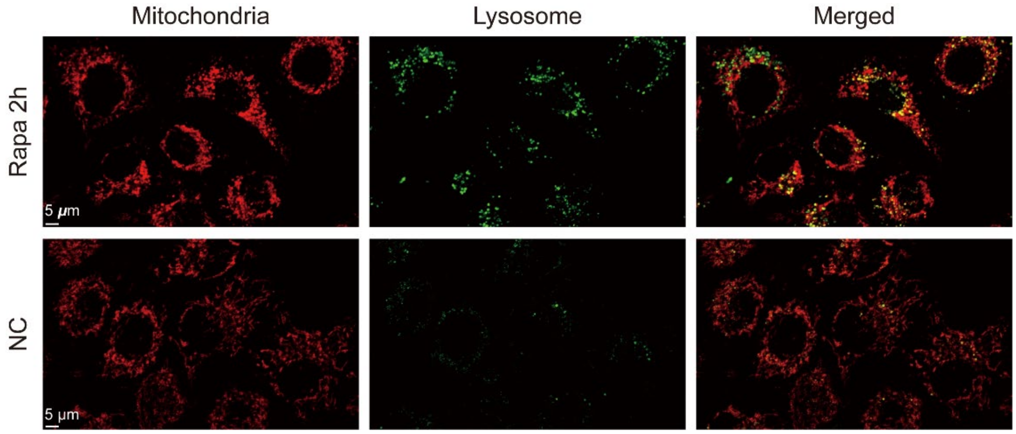

**Supplementary Figure 7. Confocal images of mitochondria and lysosome during mitophagy.** Upper panel: with rapamycin; lower panel: without rapamycin. The mitochondria were labeled with mGold-HaloTag-JF549, excited by 488 nm laser and the lysosomes were labeled with LysoTracker Deep Red, excited by 640 nm laser.

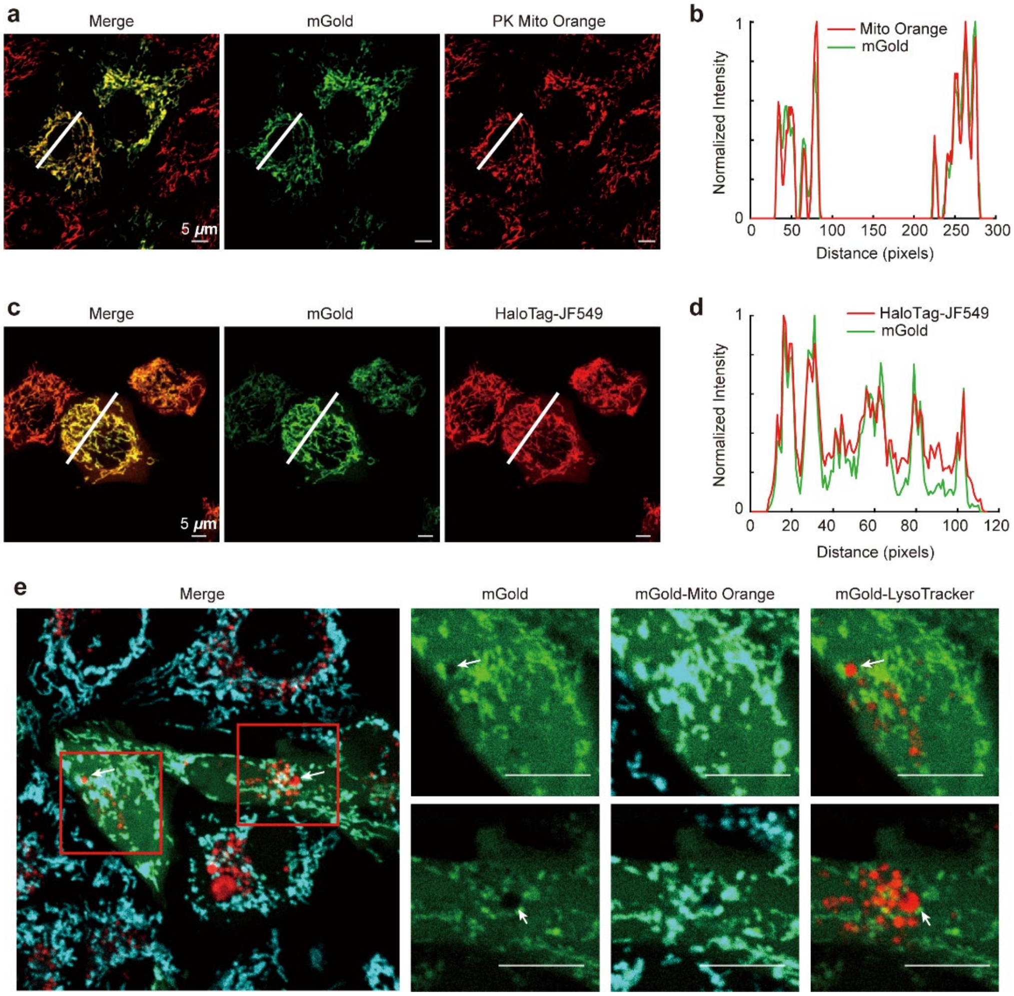

**Supplementary Figure 8. Comparison between 2×cox8-mGold-HaloTag and mitochondrial dyes.** **(a)** Confocal images of Hela cells stained with 2×cox8-mGold and commercial mitochondrial dyes, PK Mito Orange (PKMO-1, genvivotech). **(b)** Overlayed intensity distribution profile of mGold and PK Mito Orange along the white line shown in **(a)**. **(c)** Confocal images of Hela cells stained with 2×cox8-mGold-HaloTag and HaloTag dyes JF549. **(d)** Overlayed intensity distribution profile of mGold and JF549 Orange along the white line shown in **(c)**. **(e)** Confocal images of Hela cells stained with 2×cox8-mGold, PK Mito Orange, and LysoTracker Deep Red. Scale bar: 10 μm.

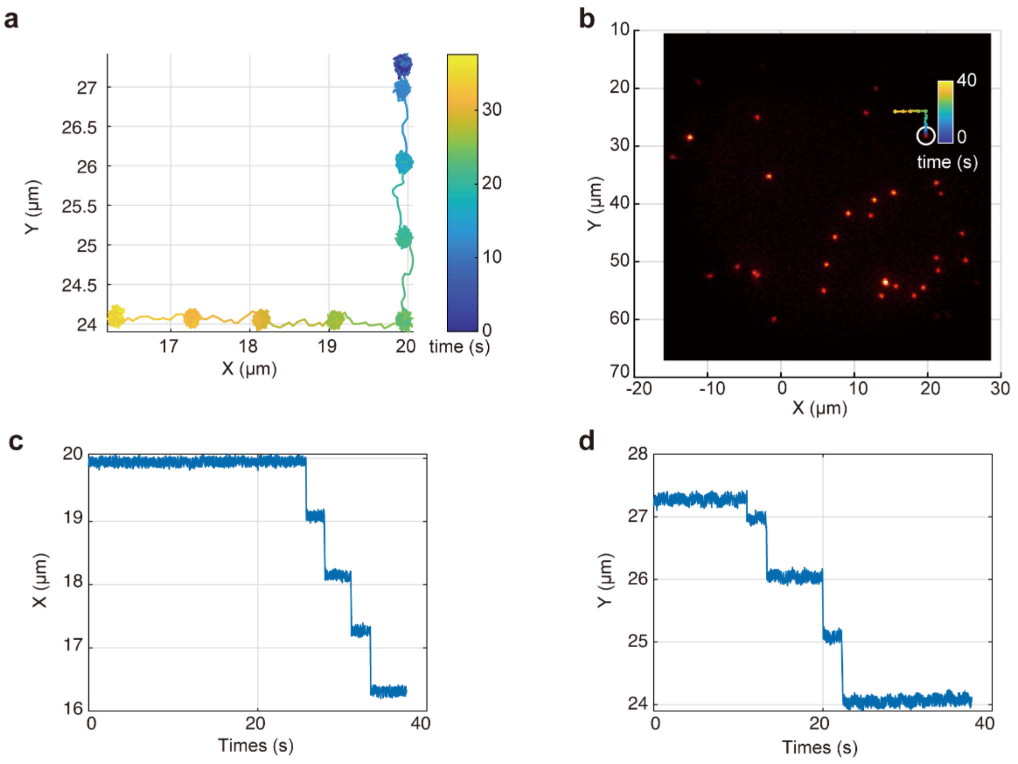

**Supplementary Figure 9. Align the tracking trajectory position with EMCCD image. (a)** Trajectories of a fluorescent bead with its position controlled by a displacement stage. **(b)** Register trajectory on EMCCD image. The white circle in the image shows the position of tracking point in the EMCCD recorded image. **(c, d)** x and y position as a function of time.

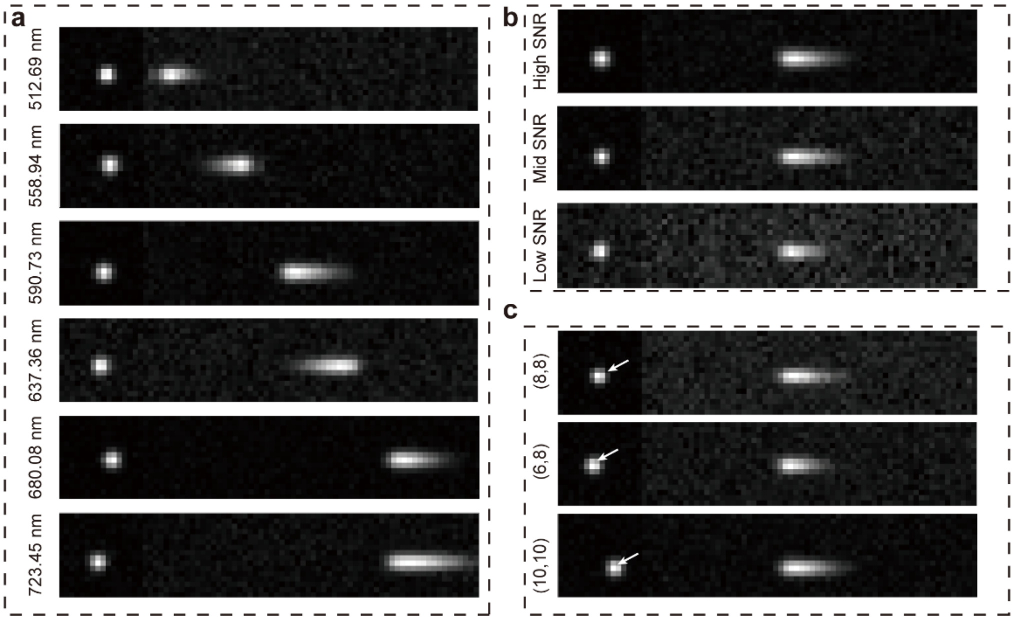

**Supplementary Figure 10. Simulated datasets for ViT model training. (a)** Simulated datasets with different spectral centroids from 510 nm to 740 nm. (b) Simulated datasets with different signal-to-noise ratio levels. (c) Simulated datasets with the particles located at different positions on image.

#### Supplementary Tables

**Supplementary Table 1.** Temporal resolution analysis of 3D-SpecDIM system.

| EMCCD parameters | | Array size of EMCCD | | |
|---|---|---|---|---|
| | | 512×512 | 25×275 | 275×25 |
| | Exposure time | **Frame rate** | | |
| 3.3 $\mu$s vertical clock speed 10 MHz pixel readout speed | 10 $\mu$s | 28 FPS | 45 FPS | 141 FPS |
| | 5 ms | 24 FPS | 37 FPS | 83 FPS |
| | 100 ms | 7.3 FPS | 8.2 FPS | 9.3 FPS |
| 0.3 $\mu$s vertical clock speed 17 MHz pixel readout speed | 10 $\mu$s | 54 FPS | 97 FPS | 644 FPS |
| | 5 ms | 43FPS | 67 FPS | 154 FPS |
| | 100 ms | 8.5 FPS | 9.1 FPS | 9.8 FPS |

**Supplementary Table 2.** Cell blebbing occurrence analysis under different conditions.

| Experiments setup | | | Observed blebbing? |
|---|---|---|---|
| Laser power (measured after objective lens) | Dyes concentration | AgNPs concentration | |
| 488 nm: 6.9 µW<br>561 nm: 0.3 - 2 µW | 1 µM NileRed for 20 min incubation | 10 µg/mL for 10 min incubation | yes |
| 488 nm: 3.5 µW | 1 µM NileRed for 20 min incubation | - | yes |
| 488 nm: 0.5 µW | 1 µM NileRed for 20 min incubation | - | no |
| 488 nm: 6.9 µW | - | 10 µg/mL for 10 min incubation | no |
| 488 nm: 6.9 µW | 1 µM NileRed for 20 min incubation | - | yes |
| 488 nm: 6.9 µW | 100 nM NileRed for 20 min incubation | - | no |
| 488 nm: 6.9 µW | - | 10 µg/mL for 10 min incubation | no |
| 488 nm: 6.9 µW | 1 µM Cellmask for 20 min incubation | 10 µg/mL for 10 min incubation | no |
| 488 nm: 6.9 µW | 1 µM DiD for 20 min incubation | 10 µg/mL for 10 min incubation | no |

**Supplementray Tables 3.** Statistics data of fluorescence spectrum centroid changes during cell blebbing.

| | Number of tracks | Average spectral changes |
|---|---|---|
| Blebbing at the periphery of cell | 5 | -7.66 ± 3.87 nm |
| Blebbing on the inner side of cell | 2 | 0.03 ± 0.24 nm |
| Without blebbing at the periphery of cell | 4 | -2.49 ± 1.30 nm |
| Without blebbing on the inner side of cell | 6 | -0.36 ± 1.87 nm |

**Supplementary Table 4.** Optical path configurations and experimental parameters.

| Experiments | Laser and Power (measured after objective lens) | F1 filter | F2 filter | F3 filter | Exposure time used on EMCCD |
|---|---|---|---|---|---|
| **Fluorescent beads spectral tracking** | Green beads, 488 nm: 0.1 μW | 535/50 (ET535/50m, Chroma) | - | - | 10 ms |
| | Yellow beads, 561 nm: 0.06 $\mu$W | 630/92(FF01-630/92, Semrock) | - | - | 10 ms |
| **Single fluorescent molecule (Setau-647) spectral tracking** | 638 nm: 2.0 $\mu$W | 630/92(FF01-630/92, Semrock) | - | - | 100 ms |
| **Mitophagy imaging** | 488 nm: 0.05-0.15 $\mu$W<br>561 nm: 0.1 $\mu$W | 488/561 (ZET488/561m, Chroma) or 405/488/561/638 (ZT405/488 | - | 630/92 or 600/52(FF01-600/52, Semrock) | 30 ms |

| | | | | | |
|---|---|---|---|---|---|
| | | /561/640mv2, Chroma) | | | |
| **Cellular membrane blebbing imaging** | 488 nm: 6.9 $\mu$W<br>561 nm: 2.0 $\mu$W | - | Polarizer (GLP10-A, Lbtek) and 488/10 (FF01-488/10, Semrock) | 630/92(FF01-630/92, Semrock) | 30 ms |

**Supplementary Table 5. Preparation of Silicon spheres coated with SLB.**

| Volume of 40 $\mu$g/mL sphere solution($\mu$L) | Volume of Tris Ca2+ buffer ($\mu$L) | Volume of lipid SUV solution ($\mu$L) | Centrifugation speed (rpm) |
|---|---|---|---|
| 1.4 | 53.6 | 125.0 | 2000 |

#### Supplementary Note

##### Supplementary Note 1: Silicon spheres coated with supported lipid bilayers.

Small unilamellar vesicles (SUVs) containing different components of lipids were prepared using sonication. Specifically, a solution mixed with 35.24 μL (25 mg/mL in chloroform) of 1,2-dipalmitoyl-sn-glycero-3-phosphocholine (DPPC, MCE; 63-89-8), and 31.28 μL (10 mg/mL in chloroform)) of cholesterol (Chol, Aladdin; C104028) or 58.4 μL DPPC only was dried under nitrogen and left in vacuum for 12 hours to completely evaporate the chloroform. Then 2 mL of Tris $Ca^{2+}$ buffer (100mM NaCl, 3mM $Ca^{2+}$, 10mM Tris, pH 7.4) was added to the dry lipid mixture and hydrated in a 55°C water bath for one hour, forming a vesicle suspension. This vesicle suspension was then put to a 4°C refrigerator for 4 hours to achieve a final concentration of 1 mM lipid mixture. The suspension was sonicated with a probe-type ultrasonicator at a constant amplitude of 60% (maximum power of 130W) for 10-15 minutes, pausing for 10 seconds after every 30-40 seconds of sonication in an ice bath to form an SUV suspension. Then, 2 mL of SUV suspension was transferred into two sterile 1.5 mL centrifuge tubes in 900 μL aliquots each and sealed with sealing film. The tubes were centrifuged at 15,000g for 1 hour at 4°C, leading to the appearance of a black substance at the bottom of the tubes (probe fragments and unbroken vesicles). For each tube, 800 μL of supernatant was collected into 1.5 mL centrifuge tubes and sealed, then stored in a 4°C refrigerator. A 100 μL aliquot of the SUV solution was then taken for particle size measurement with dynamic light scattering, resulting in 1.6 mL of 1 mM DPPC/chol or DPPC SLB solution.

The 150-nm silicon spheres (Xfnano, 7440-21-3) were diluted with Tris $Ca^{2+}$ to 40 μg/mL. According to the **Supplementary Table 5**, the spheres were further diluted to a certain concentration. The diluted microsphere solution was then heated in a 65°C water bath before being mixed with the lipid SUV solution. The mixture was maintained in the water bath for 30 minutes, with rotation every 5 minutes, and then gradually cooled to room temperature. Following the speeds indicated in the table, the mixture was centrifuged for 5 minutes. After centrifugation, the Tris $Ca^{2+}$ supernatant was replaced with Tris buffer (100 mM NaCl, 10 mM Tris, pH 7.4), resulting in 180 μL of SLB-wrapped silicon sphere solution.

For Nile Red staining, 1 μL of a 3 mM Nile Red stock solution was dissolved in 3 mL of Tris buffer to make a 1 μM Nile Red solution. Then, 5.76 μL of 1μM Nile Red was added to the SLB-wrapped silicon sphere solution (final concentration of 32 nM) and incubated in the dark at room temperature for 10 minutes.

**Supplementary Note 2: Vision Transformer and domain adaption-based spectral feature recognition for improving spectral imaging precision.**

**Vision Transformer (ViT) network**. To improve the spectral detection accuracy, we developed a spectral feature recognition method based on the Vision Transformer (ViT) model [2]. As shown in the **Extended Data Fig. 2**, the collected spectral images dimensions are 16×80 (with the left 16×16 as reference fluorescence positioning information and the right 16×64 as the dispersive spectrum, and the pixel shifting distance indicates the emission wavelengths). The ViT initiates the process by dividing an input image $I$ into a series of $N$ patches $I_p$, each reshaped into a vector of dimension $P^2 \cdot C$, where $P$ is the patch size and $C$ is the number of color channels. These vectors are then projected into a higher-dimensional space $D$ using a matrix $E \in \mathbb{R}^{(P^2 \cdot C) \times D}$, forming a sequence of patch embeddings., Then learnable positional embeddings are added to this sequence to maintain spatial information, resulting in the input of transformer encoder:

$$X_{patch} = \left[I_{p_1}E, I_{p_2}E, \dots, I_{p_N}E\right] + E_{\mathrm{pos}}, \quad (1)$$

The encoder consists of multiple layers, each comprising two main components: Multi-Head Self-Attention (MHSA) and Feed-Forward Networks (FFNs). MHSA is computed as:

$$Attention(Q, K, V) = softmax\left(\frac{QK^T}{\sqrt{d_k}}\right)V, \quad (2)$$

where $Q$, $K$, and $V$ are queries, keys, and values obtained by projecting the input patch $X_{Patch}$, and $d_k$ is the dimension of the key, ensuring proper scaling. This attention mechanism allows ViT to focus on relevant parts of the image by assigning importance weights to different patches. Following MHSA, each encoder layer applies position-wise FFNs consisting of two linear transformations with a ReLU activation. Specifically, for an input vector $x$, the FFN is:

$$FFN(x) = \max(0, xW_1 + b_1)W_2 + b_2 \quad (3)$$

Where $W_1$, $W_2$, $b_1$, and $b_2$ are learnable parameters. Layer normalization and residual connections are also employed around both MHSA and FFN modules, enhancing training stability and allowing deeper models.

By processing images through these sequential transformer layers $x' = LayerNorm(x + Sublayer(x))$, ViT captures complex patterns and dependencies among image patches, leveraging the power of the transformer architecture for spectral centroids recognition tasks. This design enables the extraction of rich feature representations and explicitly makes the model focus on spatial relationships within the image, marking a superior outcome compared to convolutional networks.

**Domain adaption strategy.** While the mentioned model demonstrates impressive performance on simulated datasets, practical applications often reveal that simulated training data (source domain) may not encompass the full range of variability encountered in the target domain. To address this challenge, we implemented a novel approach, the Regression Margin Disparity Discrepancy (RMDD), to evaluate and reduce distribution discrepancies in domain adaptation tasks (**Extended Data Fig. 2b**)[3]. In short, RMDD is a measure of the difference in distribution between the source domain and the target domain, with a particular focus on the difference in the context of margin loss. Central to this approach is the utilization of a Gradient Reversal Layer (GRL) that strategically modifies the direction of gradient flow during backpropagation, thereby encouraging the model to learn features that are invariant across domains. During the forward pass, the GRL acts as an identity function, allowing data to pass through unchanged. However, during the backward pass, it multiplies the gradient by a negative constant ($-\lambda$). This operation effectively reverses the direction of the gradient flow for the adversarial predictions, which is mathematically represented as:

$$GRL(\nabla_\theta L) = -\lambda \cdot \nabla_\theta L, \tag{3}$$

where $\nabla_\theta L$ denotes the gradient of the loss with respect to the model parameters $\theta$, and $\lambda$ is a hyperparameter that controls the strength of the gradient reversal. The RMDD loss function is defined as:

$$D_\gamma(\hat{y}_s, \hat{y}_{s_{adv}}, \hat{y}_t, \hat{y}_{t_{adv}}) = -m \cdot L_1(\hat{y}_s, \hat{y}_{s_{adv}}) + L_1(\hat{y}_t, \hat{y}_{t_{adv}}), \tag{4}$$

where $D_\gamma$ represents the disparity between the actual predictions and their adversarial counterparts in the source ($\hat{y}_s, \hat{y}_{s_{adv}}$) and target domains($\hat{y}_t, \hat{y}_{t_{adv}}$), and $L_1$ denotes the L1 loss. The term $m$ is a pre-defined constant that scales the disparity in the source domain to control its influence on the overall adaptation process.

**Loss function.** The acquisition of loss function can be separated into two stages. For the training stage, it utilizes L1 loss to train on a simulated dataset. The loss function is calculated as follows:

$$L_{reg} = \frac{1}{N}\sum_i |y_i - \hat{y}_i|, \tag{5}$$

where $N$ is the total number of observations in the dataset, $y_i$ and $\hat{y}_i$ are the actual value and predicted value of the $i$-th observation, respectively. For the inference stage, we randomly mixed practical data with simulated data and used the average spectrum of fluorescent microspheres collected under the same conditions as pseudo-labels $y_{t_{pseudo}}$ for training. Thus, the overall loss during the domain adaptation stage is the weighted sum of regression loss and RMDD loss:

$$L_{domain} = L_{reg}(y_s, \hat{y}_s) + L_{reg}\left(y_{t_{pseudo}}, \hat{y}_t\right) - D_\gamma(\hat{y}_s, \hat{y}_{s_{adv}}, \hat{y}_t, \hat{y}_{t_{adv}}), \quad (6)$$

**Training-data simulation.** To generate spectral image sequences for training, we simulated the data in different imaging parameters, as illustrated in **Supplementary Fig. 10**. The parameters include background noise, spectral centroid wavelength, diffusion coefficient, exposure time, photon count, and dichroic mirrors, et.al. Ultimately, we generated a total of 1,378,800 spectral images with varying signal-to-noise ratios, and the spectral centroids ranging from 510 to 740 nm.

**Integration of Spectral Feature Recognition Methods.** To recognize spectral features in our experiments, we utilize both a conventional fitting-based approach and a deep learning-based approach. Illustrated in **Extended Data Fig. 2**, the composite image of 16×80 pixels includes a 16×16 segment for molecular position and a 16×64 segment for spectral data after dispersion, with both segments undergoing normalization. Initially, the molecular coordinate $p$ within the 16x16 segment is located. Aligning with the practical optical path configuration, we designate three spectral windows within the spectral image for 488 nm (from pixel 1 to pixel 22), 561 nm (from pixel 23 to pixel 43), and 640 nm (from pixel 44 to pixel 64) emissions. The channel with the largest signal is selected for normal distribution fitting to identify the pixel coordinates $s$ of the spectral centroid. The discrepancy in pixels between the spectral centroid $s$ and positional data $p$ is calculated and converted into the actual wavelength of the spectral emission peak via a calibration function[4]. Alternatively, the neural network method inputs spliced images into a pre-trained model, which then outputs the spectral predictions directly.

**Training procedure and Testing details.** Totally 1,378,800 images across a range of signal-to-noise ratios were generated for the training of our deep learning model, reserving 10% of these for testing purposes. For domain adaptation, we collected tracks from fluorescent beads at 9 distinct signal-to-noise ratios to fine-tune the model. We further assessed the model's versatility by employing various spectral window settings. The training was conducted using two NVIDIA GeForce RTX 3090 graphics cards, with the programming carried out in the PyTorch-Lightning framework. Optimization was managed by the Adam optimizer, and we processed the data in batches of 128. Detailed information about our model is available on our GitHub page.

**Supplementary Notes 3. Multi-resolution imaging with 3D-SpecDIM.**

To reconstruct a 3D volumetric image of the mitophagy process, we employed two avalanche photodiodes (APDs) to simultaneously collect signals from mitochondria and lysosomes. Specifically, a 600/50 filter (FF01-600/52, Semrock) was positioned in front of APD1 (as shown in **Extended Data Fig. 1**) to enable active tracking based on HaloTag-JF549 signals. A 706/95 filter (ET706/95, Chroma) was placed in front of APD2 to capture lysosomal signals. The photon addressable capability of 3D-SpeDIM facilitates us to reconstruct the 3D image that surrounding target particle by registering the fluorescence photons to laser focus positions in a separated detection channel. Here the 3D image of lysosome was reconstructed in APD2. The recorded data included EOD scanning coordinates (XY information) and TAG phase (Z information) corresponding to the arrival of each photon. During data processing, we binned the photon counts at 100 ms intervals and allocated them into a 3D data voxel of dimensions 5×5×10, based on their coordinate values. When there is an overlap of EOD and TAG scanning voxel units at different trajectory points, we use the value with the higher photon count as the reconstruction vaule. Finally, by integrating these 3D data cubes with the 3D tracking trajectory, the 3D distribution of lysosomal morphology along the trajectory can be reconstructed. The 3D visualization of multi-resolution image was performed with Avizon software.
